# Cell-type-specific gene expression and regulation in the cerebral cortex and kidney of atypical *Setbp1*^S858R^ Schinzel Giedion Syndrome mice

**DOI:** 10.1101/2023.07.31.551338

**Authors:** Jordan H. Whitlock, Tabea M. Soelter, Timothy C. Howton, Elizabeth J. Wilk, Vishal H. Oza, Brittany N. Lasseigne

## Abstract

Schinzel Giedion Syndrome (SGS) is an ultra-rare autosomal dominant Mendelian disease presenting with abnormalities spanning multiple organ systems. The most notable phenotypes involve severe developmental delay, progressive brain atrophy, and drug-resistant seizures. SGS is caused by spontaneous variants in *SETBP1*, which encodes for the epigenetic hub SETBP1 transcription factor (TF). *SETBP1* variants causing classical SGS cluster at the degron, disrupting SETBP1 protein degradation resulting in toxic accumulation, while those located outside cause milder atypical SGS. Due to the multi-system phenotype, we evaluated gene expression and regulatory programs altered in atypical SGS by snRNA-seq of cerebral cortex and kidney of *Setbp1*^S858R^ heterozygous mice (corresponds to the human likely pathogenic *SETBP1*^S867R^ variant) compared to matched wild-type mice by constructing cell-type-specific regulatory networks. *Setbp1* was differentially expressed in excitatory neurons, but known SETBP1 targets were differentially expressed and regulated in many cell types. Our findings suggest molecular drivers underlying neurodevelopmental phenotypes in classical SGS also drive atypical SGS, persist after birth, and are present in the kidney. Our results indicate SETBP1’s role as an epigenetic hub leads to cell-type-specific differences in TF activity, gene targeting, and regulatory rewiring. This research provides a framework for investigating cell-type-specific variant impact on gene expression and regulation.

## Introduction

Schinzel Giedion Syndrome (SGS) is an ultra-rare autosomal dominant Mendelian disease caused by spontaneous variants in *SETBP1*.^1,2^ SGS patients, who do not typically survive past infancy, present with gastrointestinal, cardiorespiratory, neurological, musculoskeletal, and urogenital abnormalities as well as cancer. However, the most notable phenotypes involve severe developmental delay, progressive brain atrophy, and drug-resistant seizures.^2–4^ *SETBP1* encodes for the SETBP1 transcription factor (TF), which acts as an epigenetic hub and has many understudied functions in SGS.^5^ *SETBP1* variants associated with classical SGS cluster at mutational hotspots in the degron lead to disruptions in protein degradation resulting in a toxic accumulation of SETBP1 protein, therefore making SGS a gain-of-function (GoF) disorder.

Alternatively, variants outside the degron, reported in cases of atypical SGS, exhibit a milder form of the syndrome and longer lifespan, classified by microcephaly, facial gestalt, cardiac defects, alacrima, hypertonia, developmental delay, seizures, spasticity, and vision impairment.^6–8^

Prior human cell and mouse model studies uncovered multiple potential mechanisms altering neurodevelopment in SGS (reviewed in^9^), documenting the role of SETBP1 accumulation in inhibiting p53 function, causing aberrant proliferation, deregulating oncogenes and suppressors, and promoting unresolved DNA damage and apoptosis resistance in neural progenitor cells (NPCs).^10^ *SETBP1* variants were also implicated in decreased genome pan-acetylation.^4^ Furthermore, variants resulting in SETBP1-mediated PP2A inhibition have been found somatically in several myeloid malignancies as well as in germline variants related to altered neurodevelopment.^9^ SGS’s syndromic presentation and SETBP1’s TF role provide the opportunity to study the impact of a variant on cell-type-specific expression and regulation in a context-specific framework.

However, no studies to date have determined if these molecular alterations are present in other affected cell types or tissues, like the kidney, where hydronephrosis and recurrent urinary tract infections are secondary phenotypes of SGS.

Multiple biological processes have been hypothesized to explain tissue-specific manifestations of disease phenotypes.^11^ To understand the multi-system phenotype of SGS, we evaluated gene expression and regulatory programs altered in atypical SGS through single-nuclei RNA-sequencing (snRNA-seq) of the cerebral cortex and kidney of 6-week-old male *Setbp1*^S858R^ heterozygous mice compared to age- and sex-matched wild-type (WT) mice and construction of cell-type-specific regulatory networks **(Fig 1)**. The *Setbp1*^S858R^ variant corresponds to the human likely pathogenic *SETBP1*^S867R^ atypical SGS variant, located within the SKI homologous domain (amino acids 706 - 917), a region named for its partial homology to the SKI oncoprotein.^12^ In addition to determining cell-type-specific impacts of the *Setbp1*^S858R^ variant in two SGS-affected tissues, our research provides a framework for investigating changes introduced by a variant to mechanisms of cell-type-specific expression and regulation in this understudied rare disease.

**Figure 1.**
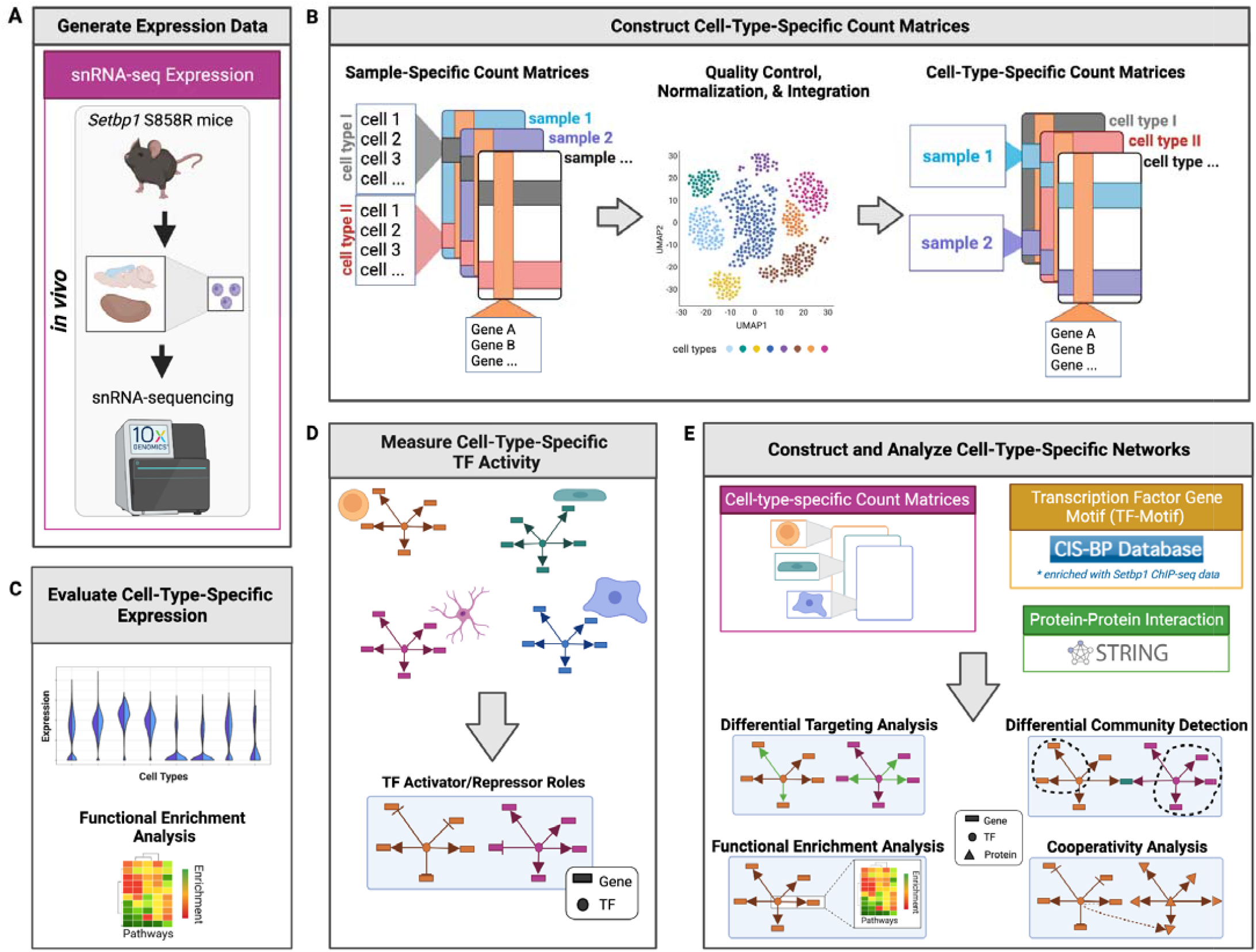
Schematic overview of our approach (A) We generated single-nuclei RNA-seq (snRNA-seq) from male C57BL6/J (WT) and *Setbp1*^S858R^ cerebral cortex and kidney tissues. **(B)** We processed and aggregated our data across samples to create cell-type-specific count matrices. **(C)** Next, we assessed cell-type-specific expression for *Setbp1* and genes SETBP1 is known to target. **(D)** With decoupleR, we measured cell-type-specific TF activity. **(E)** We acquired protein-protein interaction data (PPI; green) from STRING and Transcription Factor-motif data (TF-motif; yellow) from CIS-BP enriched for SETBP1 targets. We then built cell-type-specific TF-gene regulatory networks with the data we generated by using the message-passing algorithm PANDA to identify regulatory relationships between TFs (circles), genes (rectangles), and proteins (triangles) for each S858R and WT cell type in both tissues and investigated differential communities using ALPACA, differential gene targeting, cooperativity analysis, and functional enrichment analysis.

## Materials and methods

A more detailed version of all data processing and analysis steps is available in the **Supplemental Methods**.

### *In-vivo* Mouse Samples and Nuclei Isolation

We obtained decapsulated kidneys and cerebral cortex hemispheres from three 6-week-old male C57BL/6J-*Setbp1*^*em2Lutzy*^/J mice heterozygous for *Setbp1*^S858R^ (JAX Stock #033235) and three wild-type (WT) age and sex-matched C57BL6/J mice (JAX Stock #000664) **(Table S1)**.^13^

We adapted and conducted single nuclei isolation for the kidney as previously described.^14^ We resuspended the final nuclei pellet in Nuclei Suspension Buffer with a final yield of 1.3 - 3.0 million nuclei per kidney. We adapted our single nuclei isolation for cerebral cortex samples for mice from the previously described^15^ and sorted nuclei using fluorescence-activated cell sorting (FACS) on an Atlas BD ARIA II with a yield of 1.5 - 3.0 million nuclei per cerebral cortex hemisphere.

### snRNA-seq Processing

We generated single-nuclei RNA-sequencing (snRNA-Seq) libraries using the 10X Chromium Single Cell 3_LJ_ v3 (10X#: PN-1000121) protocol with a targeted cell recovery rate of 20,000 nuclei per sample. The libraries were indexed and sequenced on a NovaSeq 6000 (Illumina).

We processed all snRNA-seq data using 10X Genomics Cell Ranger (v6.1.1). We aligned sequencing reads to *mus musculus* (mm10). We confirmed all variants post-sequencing through conversion of the .bam output files from Cell Ranger to .vcf using bcftools (v.1.6) and visualizing the loci using the Integrative Genome Viewer (IGV) 2.8.0.^16,17^ The S858R variant was present in all heterozygous samples at chr18:78857841-78857920 **(Figure S1)**. We mapped the mouse S858R variant to the human Schinzel Gideon-causing variant, *SETBP1* S867R, using ConVarT (accessed February 2022).^18^

### Setbp1 Protein Quantification

We used the Mouse SETBP1 ELISA kit (MyBioSource Cat No. MBS9335445) to quantify SETBP1 protein abundance in kidney and cerebral cortex samples for both conditions (n = 3 replicates per condition per tissue). We performed incubation and manual washing as described in the MyBioSource Kit protocol and measured absorbance using a Tecan Infinite M Plex plate reader at 450 nm for 10 flashes. The proportion of total protein abundance in □g/mL was measured in all tissues (n = 3 per condition per tissue) using the BCA with the Pierce protocol.^19^ Significance was calculated using a paired t.test between groups (p-value < 0.05) **(Figure S2)**.

### Detection of differentially expressed genes

We applied the Seurat FindAllMarkers function with an avg_log2FC threshold of 0.1. Results for all DEGs (p-adjusted value < 0.05) for all S858R cerebral cortex and kidney cell types (**cell type markers; Table S2)** are in **Tables S3, S4**. Using gProfiler2 (v0.2.1), we identified significantly enriched pathways of up and down-regulated differentially expressed genes from Gene Ontology (GO; 2022-03-22 release), Reactome (REAC; 2022-5-25 release), KEGG (2022-05-26 release), TF (TRANSFAC Release 2021.3), MIRNA release 7.0, HPA (21-12-17 release), CORUM 3.0, and HP (hp.annotations 12) sources^20^. We performed functional enrichment analysis (FEA) on the DEGs filtered for the SETBP1 target gene set (gene set construction**; Supplemental Methods, Table S5)** and applied a p-adjusted threshold of 0.05 and a Bonferroni procedure for multiple hypothesis correction.

### Gene set enrichment analysis (GSEA)

We performed GSEA with VISION (v3.0.1)^21^ using mouse gene sets from the Molecular Signature Database (MSigDB) (v2023.1.Mm) for hallmarks, Polycystic kidney disease, and PP2A phosphatase (GO:0051721).^22–24^ We calculated Polycystic kidney disease and PP2A phosphatase signatures using Seurat’s AddModuleScore.

### TF Activity

We inferred transcription factor (TF) activity using decoupleR^25^ (v.2.6.0) by combining our scRNA-seq data with prior knowledge from CollecTRI, a comprehensive network of TFs and their direction of regulation on transcriptional targets (accessed May 2023).^26^ We calculated activity scores for all 732 TFs by using a Multivariate Linear Model, including TFs with a minimum of 5 targets. We summarized average TF activity by each cell type. Then we scaled and centered data to prioritize TFs with the largest change in activity between conditions using absolute percentage change of the average activity. More positive percent changes (above Q3) represented a larger magnitude of TF activity in the S858R, and the more negative percent changes (below Q1) represented a larger magnitude of TF activity in WT.

### Cell-type-specific Network Construction and Analysis

To model cell-type-specific regulatory relationships between TF and genes within both conditions, we constructed cell-type-specific networks using Passing Attributes between Networks for Data Assimilation (PANDA)(v.1.2.1).^27,28^ Network edge weights represent the strength in interactions between nodes (co-expressed genes, TFs, or pairs of TFs and genes) where more positive weights correspond to higher confidence interactions in a given cell-type-specific network. All analysis was performed in a High-Performance Computing environment at UAB.

Gene targeting enabled us to quantify all in-degree edge weights of each gene. A higher targeting score indicated a gene had a higher sum of all in-degree edge weights. By calculating a differential targeting score for a gene between two networks, we could compare if a gene is targeted more or more “important” in one network versus the other. We calculated differential gene targeting between conditions by taking the difference in targeting scores for each gene, with positive scores indicating enrichment in S858R mice. Genes with differential targeting scores above the third quartile were selected within each cell type and further investigated using GSEA after filtering for the SETBP1 gene set (p-adjusted < 0.05).

We conducted a comparative analysis of regulatory network community structures in all cell types for both conditions by employing the bipartite community detection algorithm, COmplex Network Description Of Regulators (CONDOR)(v.1.2.1)^29^, on positive edge weights from the PANDA networks. We used ALtered Partitions Across Community Architectures (ALPACA)(v.1.2.1)^30^, a message passing and modularity optimization method, to detect structural changes in closely interacting groups of nodes between conditions referred to as differential communities and quantified the similarities between SETBP1 communities across cell types within each tissue using the Jaccard similarity index (JI).

To understand regulatory network rewiring due to the *Setbp1*^S858R^ variant, we filtered regulatory (TF-gene) and cooperativity (protein-protein) networks to only include edges associated with the SETBP1 gene set. We then annotated the regulatory and cooperativity network edges, categorizing them based on the presence or absence of regulation and cooperation between genes and proteins, and investigated specific proteins involved in mechanisms related to altered cell cycle chromatin remodeling, DNA damage, and phosphorylation in SGS.^9^

### SETBP1 targets are expressed in a cell-type-specific manner in atypical SGS *Setbp1*^**S858R**^ mouse cerebral cortex and kidney tissues

As prior studies using patient-derived cell lines and cerebral organoids indicated that variants in *SETBP1* lead to an accumulation of SETBP1 protein in SGS,^10^ we confirmed increased SETBP1 protein abundance in the cerebral cortex and kidney S858R tissues compared to WT (p-value < 0.05, paired t-test) (**Figure S2**). After profiling single-nuclei RNA from the cerebral cortex (n = 51,038 total nuclei) and kidney (n = 78,751 total nuclei), we confirmed the presence of the variant in all S858R samples (**Figure S1**) and assigned cell types (**Table S2**, and **Figures S3, S4**). We identified differentially expressed genes (p-adjusted < 0.05 Wilcoxon rank sum) between S858R and WT and found widespread gene expression changes impacting all cell types in our study (**Tables S3, S4)**. *Setbp1* was ubiquitously expressed across all cell types in both conditions, in line with prior studies **(Figure 2A, 2B)**.^31^ However, *Setbp1* was only significantly differentially expressed (upregulated) in S858R excitatory neurons compared to WT (p-adjusted < 0.05, Wilcoxon rank sum) (**Fig 2C**). Functional enrichment analysis (FEA) on all statistically significant differentially expressed genes in atypical SGS mice revealed cell-type-specificity in the cerebral cortex. These genes were overrepresented in pathways related to previously noted SGS signatures and clinical phenotypes such as cell death, apoptosis language impairment, and neurodegeneration (**Figures S5-S10)**. There was no enrichment for differentially expressed genes in kidney cell types associated with polycystic kidney disease to support a cystic phenotype in SGS hydronephrosis^32^ or in genes related to failed-repair, a common kidney disease mechanism^33^ **(Figures S11, S12, Supplemental Methods)**. DNA damage accumulation has been previously confirmed in NPCs for SGS; however, to our knowledge, it has not been studied within the context of SGS kidneys. Interestingly, we found kidney cell-type-specific decreases in DNA repair, WNT-Beta Catenin signaling, and p53 pathway expression signatures in atypical SGS mouse kidneys (**Figure S13B)**. Due to SETBP1’s role as a TF, we investigated mouse orthologs of known SETBP1 targets for cell-type-specific differential expression between S858R and WT **(Fig 2C, 2D)**. We found that many target genes with functions in SGS-associated pathways were differentially expressed (p-adjusted < 0.05, Wilcoxon rank sum) **(Table 1)**.

**Table 1.**
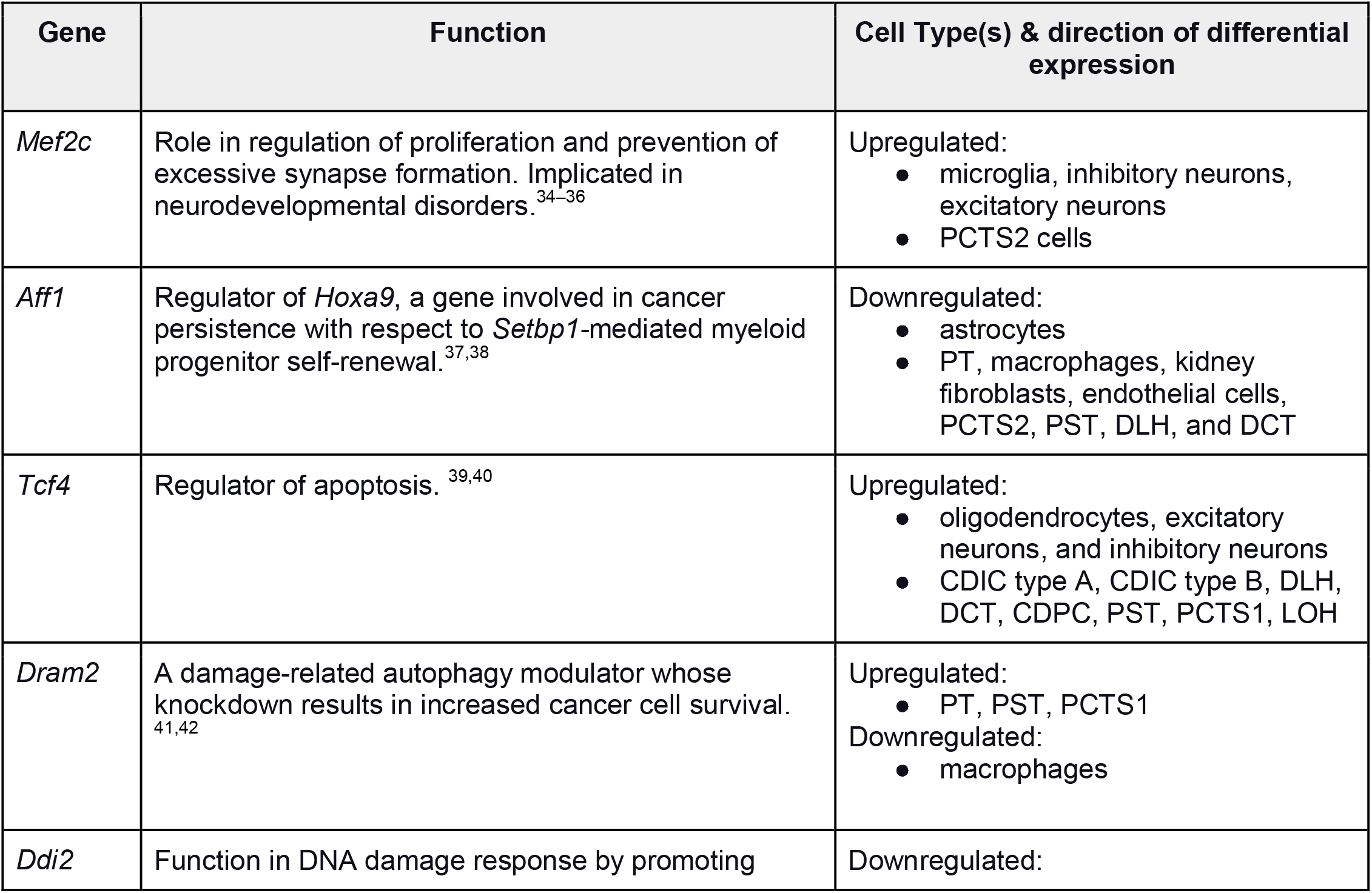
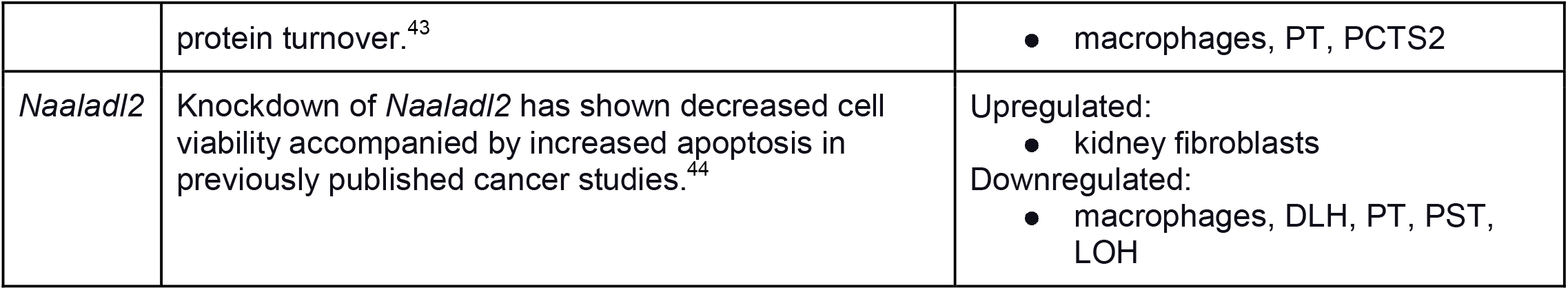
Highlighted SGS-signature associated differentially expressed SETBP1 targets in S858R cerebral cortex and kidney cells.

**Figure 2:**
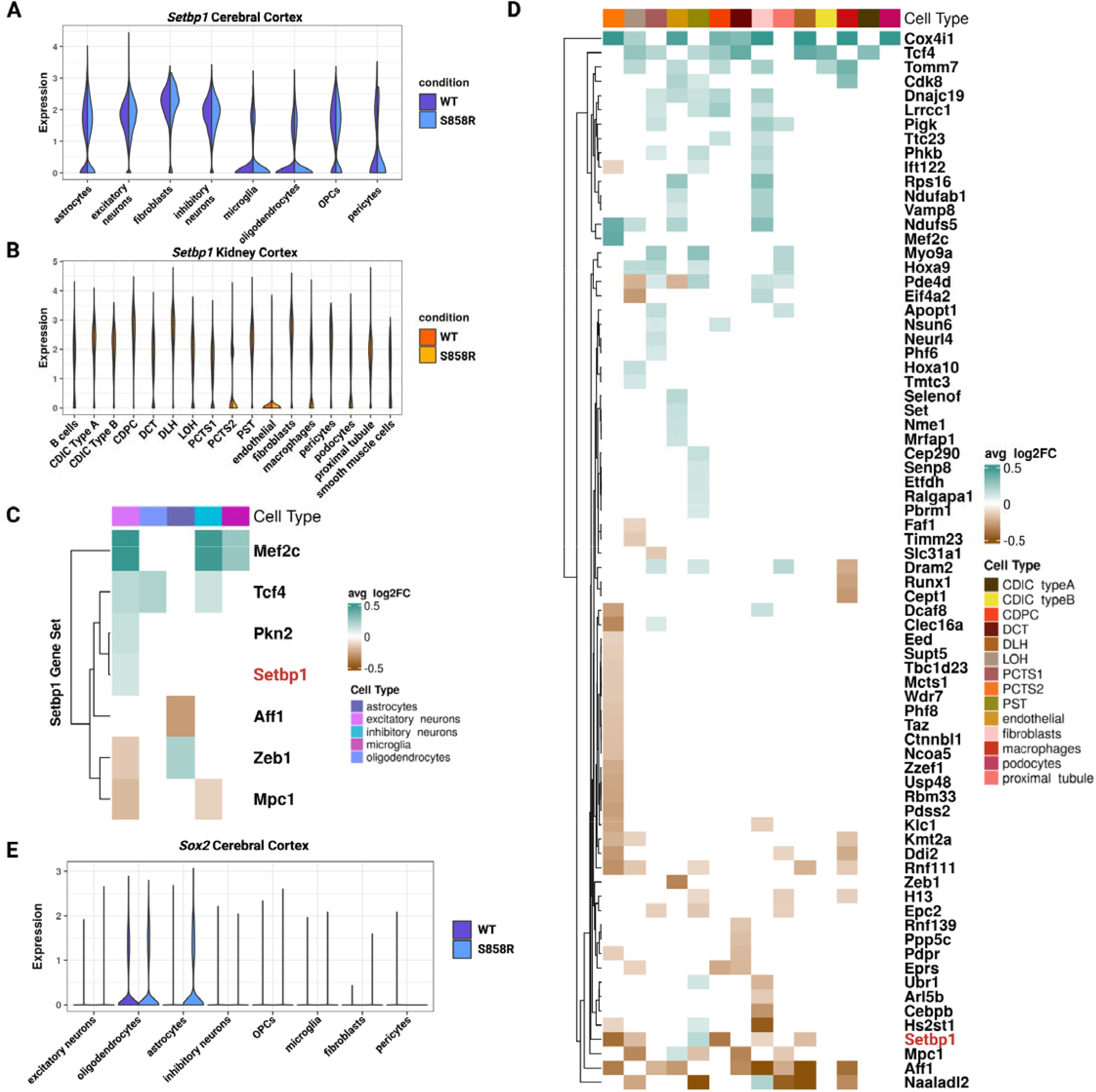
Differential expression of SETBP1 targets reveals cell-type-specific signatures of apoptosis resistance, oncogene disruption, and altered cell cycling in NPC-derived cells: Expression of *Setbp1* in S858R and WT **(A)** cerebral cortex and **(B)** kidney cells. Differentially expressed known SETBP1 target genes (avg_log2FC threshold of 0.1 and p-adjusted value < 0.05) in S858R and WT **(C)** cerebral cortex and **(D)** kidney cells. Teal and brown denote up and downregulated genes, respectively. (**E**) Split violin plot of the expression of *Sox2* (a marker of proliferative NPCs) in S858R and WT cerebral cortex cell types.

Because prior work in SGS NPCs has indicated SETBP1 accumulation leads to a neurodegenerative phenotype, inducing p53 inhibition and genotoxic stress accompanied by altered cell cycle and DNA damage^10^, we hypothesized S858R neurons and astrocytes (cell types derived from NPCs^45^) and macrophages like microglia (previously implicated in inflammatory processes in other neurodegenerative disorders^46^) may also exhibit these molecular phenotypes. We determined the S858R oligodendrocytes, oligodendrocyte precursor cells (OPCs), and excitatory neurons exhibited a decreased enrichment score for the p53 pathway compared to WT. We also found S858R cortical pericytes had reduced DNA repair, p53, and apoptosis, while microglia, OPCs, and excitatory neurons had decreased G2M enrichment compared to WT. Additionally, S858R excitatory neurons were enriched for apoptosis compared to WT **(Figure S13A)**. Due to the presence of these SGS-associated signatures, we profiled the expression of the proliferative NPC marker *Sox2* and found exclusive expression in S858R astrocytes, suggesting they may have a disease-specific proliferative NPC population (**Fig 2E, Supplemental Methods)**.^10,47^ While previous research has indicated inflammatory conditions or disease states such as Alzheimer’s disease^48^ drive mature astrocytes to re-acquire NPC-like properties,^49^ we did not find differences in gene expression of the reactive astrocyte marker *Gfap* in astrocytes between conditions **(Figure S14, Supplemental Methods)**.

Accumulation of SETBP1 is known to result in the stabilization of SET and inhibition of PP2A in SGS.^10^ We found ubiquitous expression of *Set* in all cell types and tissues **(Figure S15, Supplemental Methods)**; however, *Ppp2ca* (the mouse ortholog for *PP2A*) was expressed in WT but not S858R astrocytes and S858R but not WT microglia **(Figure S16, Supplemental Methods)**. There was no difference in other cell types between the S858R and WT cells of both tissues. PP2A’s tumor suppressor function is known to be disrupted in SGS, and a lack of apoptosis in astrocytes has previously been noted as well.^10,9^ As reported above, our gene set signature enrichment analysis revealed a decrease in the G2M checkpoint signature in S858R astrocytes compared to WT, suggesting there may be more astrocytes in the G0/G1 cell cycle phase (**Figure S13A**). This is further supported by a subsequent downregulation of cell death and apoptotic processes in the pathways associated with downregulated SETBP1 target genes in S858R astrocytes (**Figure S13A)**.

### SETBP1 and known targets’ TF activity is altered across cerebral cortex and kidney cell types

Due to cell-type-specific changes in the gene expression of known SETBP1 targets, we inferred the TF activity of *Setbp1* and its TF targets in the cerebral cortex and kidney using a multivariate linear model.^25^ We prioritized transcriptional regulators displaying the highest absolute percent change between WT and S585R and found that SETBP1 and its targets display cell-type- and disease-specific alterations in TF activity in the cerebral cortex and kidney (**Fig 3**). More specifically, in the cerebral cortex (**Fig 3A**), SETBP1 is upregulating its targets in WT pericytes and microglia. However, SETBP1 functions as a transcriptional repressor in S858R astrocytes. Interestingly, *Setbp1* itself was not differentially expressed in these cell types. In the kidneys (**Fig 3B**), SETBP1 serves as a repressor in WT Loop of Henle (LOH), Smooth muscle cells (Smcs), B cells, Collecting Duct Principal Cells (CDPC), and Distal Convoluted Tubule (DCT) and an activator in pericytes. On the other hand, in the S585R variant, SETBP1 had a transcriptional activator role in Distal Loop of Henle (DLH) and Proximal Convoluted Tubule Segment 2 (PCTS2) cells. Similar to gene expression, the TF activity changes for SETBP1 due to the S585R variant in the cerebral cortex and kidney are subtle. However, we see additional changes in the TF activity of transcriptional regulators targeted by SETBP1.

**Figure 3:**
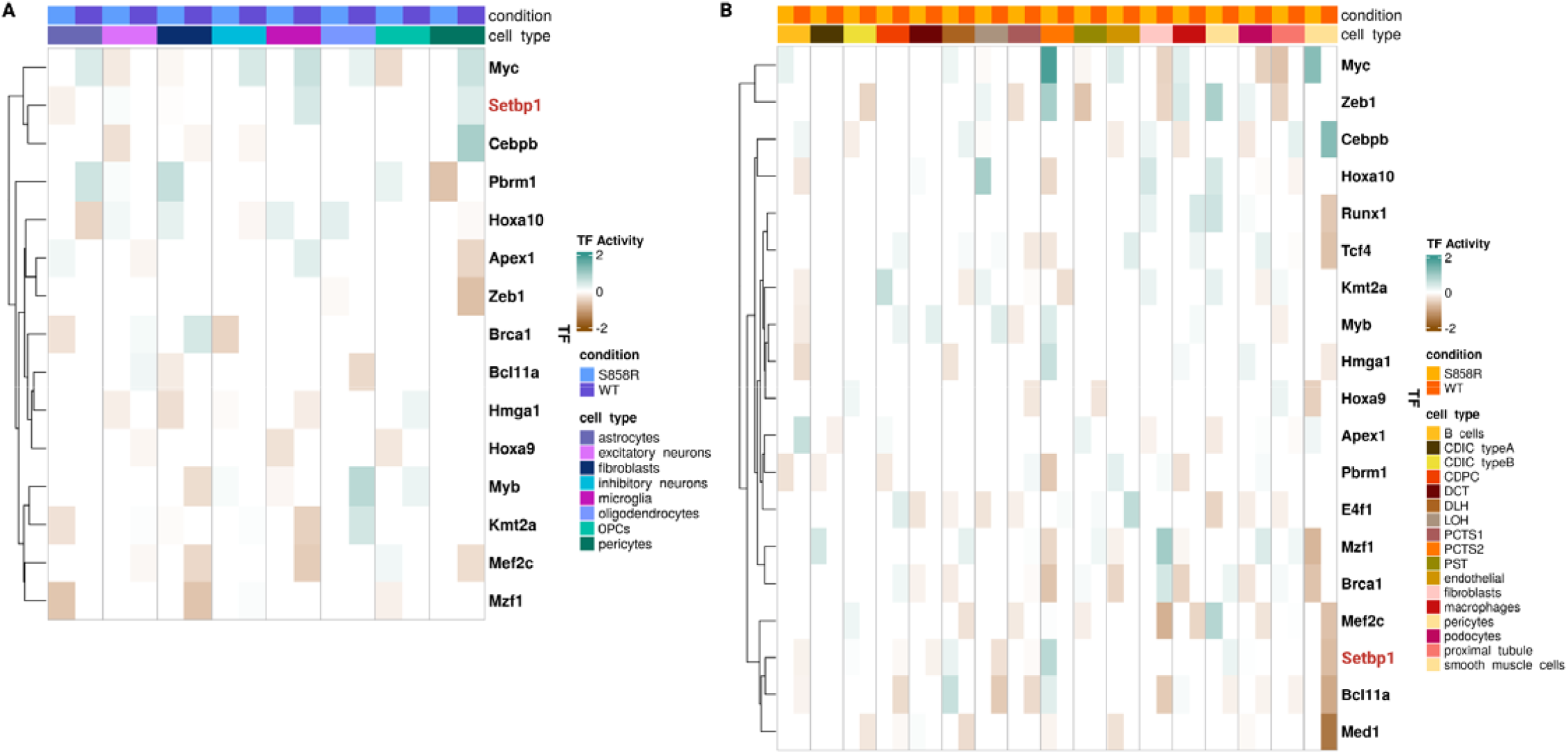
SETBP1 TF activity is altered in pericytes and microglia and multiple kidney cell types of S858R mice: Heatmap of decoupleR TF activity scores of SETBP1 and its target genes that displayed the largest percent change in activity in the cerebral cortex (**A**) and kidney **(B)**. Teal and brown correspond to increased and decreased TF activity, respectively.

The TFs regulated by Setbp1 display cell-type- and disease-specific TF activity changes. We quantified the percent change of all TFs between WT and S585R and then subsetted for those that SETBP1 transcriptionally regulates. We see 14 of these transcriptional regulators in the cerebral cortex (**Fig 3A**).

Interestingly, 13 of the 14 TFs were not differentially expressed. *Mef2c* was differentially expressed in excitatory and inhibitory neurons and microglia. We see the largest change in the TF activity of *Mef2c* in microglia and no change in TF activity in inhibitory neurons. In the kidney, we see altered TF activity in SETBP1 and 18 of its targets. These TF activity changes occur in every kidney cell type (**Fig 3B)**. Although subtle, these results indicate that the S585R variant alters the gene regulatory landscape not only through differential gene expression of direct targets but also by altering the activity of additional transcriptional regulators.

### Cell-type-specific changes in gene targeting recapitulate SGS molecular features in many atypical SGS

#### *Setbp1*^**S858R**^ mouse cell types

We constructed TF-gene regulatory networks for each cell type by condition using the message-passing algorithm PANDA^27^ to calculate the ‘responsibility’ of a TF for regulating a gene and the ‘availability’ of that gene for regulation by that TF. We then averaged the responsibility and availability networks to obtain a consensus gene regulatory network per cell type. From these networks, we calculated the normalized differential targeting score for all genes by cell type in the cerebral cortex (**Fig 4A)** and kidney (**Fig 4B**). This revealed higher total differential targeting scores in specific cell types of S858R compared to WT for both tissues, suggesting increased gene regulation, consistent with gain of function, in the S858R (**Table S7**). We performed FEA on both tissues’ differentially targeted genes from the SETBP1 gene set. In the cerebral cortex, microglia and excitatory and inhibitory neurons had differential expression of pathways indicating altered H3K9 acetylation and regulation in S858R compared to WT (**Fig 4C**). These findings are in line with recent studies that show SET overexpression alters histone acetylation and accumulation of SET can impair chromatin accessibility at distal regions of the genome, specifically, chromatin loop contacts in SGS NPCs.^4^ Our FEA also identified sloping forehead as a phenotype term associated with the excitatory neuron S858R gene signature, concordant with clinical indications that individuals with SGS have pronounced craniofacial features (i.e., excessive posterior sloping of the forehead).^50^ In the kidney, we also identified altered H3K9 signatures in S858R Collecting Duct Intercalated Cells (CDIC) type A and B and pericytes. Furthermore, S858R LOH, DLH, Proximal Convoluted Tubule Segment 1 (PCTS1), and DCT cells displayed differential pathway enrichment for cellular response to lectin, previously implicated in renal injury (**Fig 4D**).^51,52^ Overall these results demonstrate that SETBP1 with the S858R variant differentially targets genes by cell type and that molecular programs associated with typical SGS are reflected in the cerebral cortex and kidney of this model of atypical SGS.

**Figure 4:**
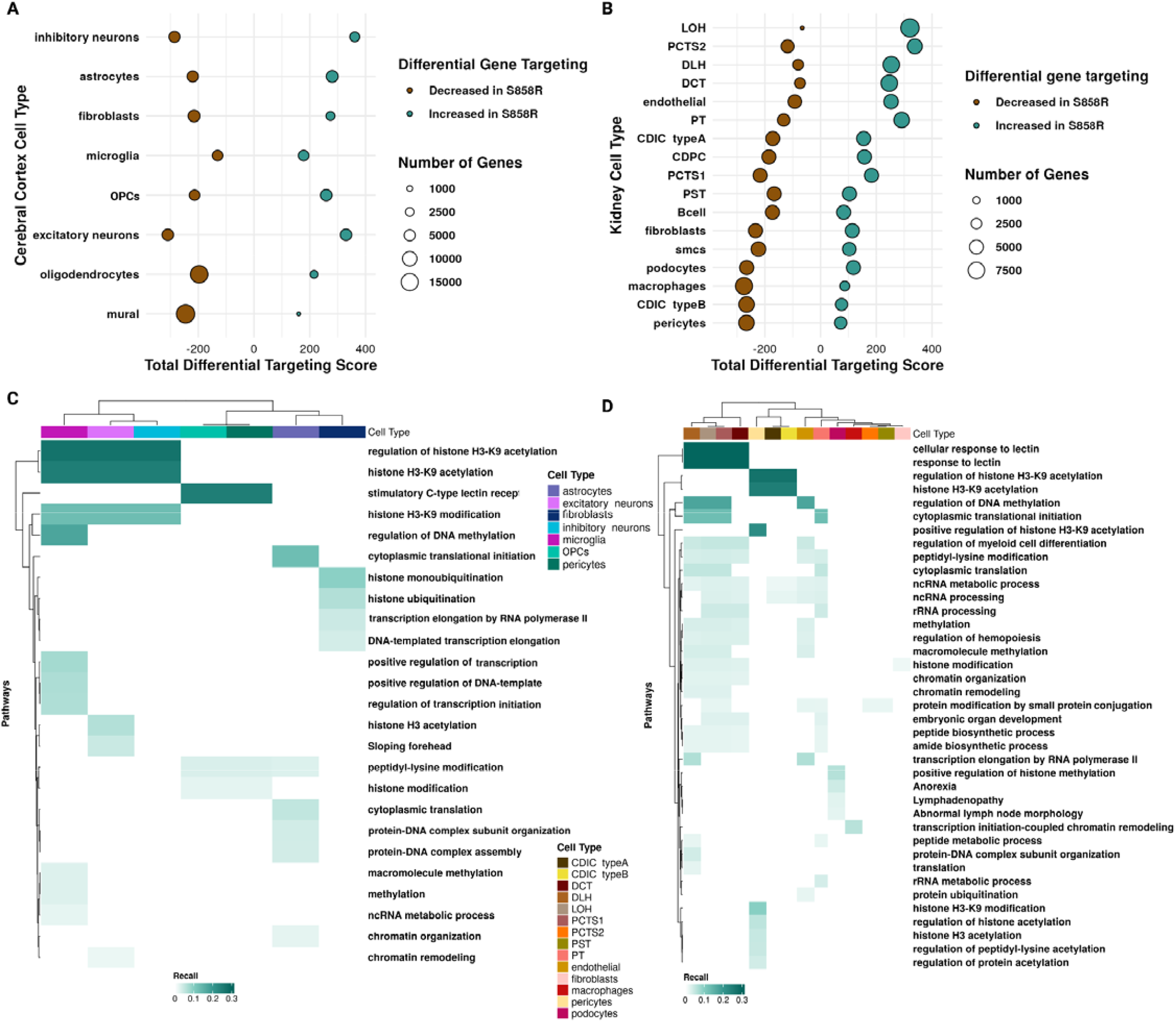
Increased gene targeting in S858R cerebral cortex and kidney cells indicates increased gene regulation and altered cell-type-specific mechanisms associated with SGS: Dot plot of the normalized total differential gene targeting by **(C)** cerebral cortex and **(D)** kidney cell types (teal for increased gene targeting in S858R compared to WT and brown for decreased gene targeting in S858R compared to WT; dot size corresponds to gene number). FEA of differentially targeted SETBP1 gene set in **(C)** cerebral cortex and **(D)** kidney cell types.

### Differential communities between S858R and WT containing SETBP1 indicate shared SETBP1 regulatory patterns between sets of cell types

We identified communities in each cell-type-specific network based on differential modularity (a metric measuring the extent to which edges in communities of S858R networks differ from those of reference WT networks) for both the cerebral cortex and kidney using the ALPACA framework.^30^ When we analyzed differential communities containing SETBP1 as a TF, we found that excitatory neurons and astrocytes had the highest Jaccard similarity index (JI) (0.42, **Fig 5A**) and were also more similar to OPCs (0.44 and 0.38, respectively) than to other cerebral cortex cell types. Oligodendrocytes and pericytes, not previously suggested to have a role contributing to SGS, had a JI of 0.59, concordant with having similar pathway signatures in S858R compared to WT as described above (**Figure S17**). These two cell types also had high JI with fibroblasts (0.41 for both). In kidney, we found SETBP1-specific differential communities were most similar between DCT, DLH, PCTS2, Proximal Tubule (PT), endothelial, and LOH cell types (JI = 0.30 to 0.53), between fibroblasts, macrophages, pericytes, podocytes, and Smcs (JI = 0.38 to 0.64), and between B cells, Proximal Straight Tubule (PST), CDIC type A and B, and CDPC (JI = 0.34 to 0.51) (**Fig 5B**). Overall, we found that there are sets of cell types with more similar SETBP1 differential communities, further underscoring the multifunctional role of SETBP1 in atypical SGS and that additional cell types are impacted by, may contribute to, and work together in disease etiology and pathogenesis.

**Figure 5:**
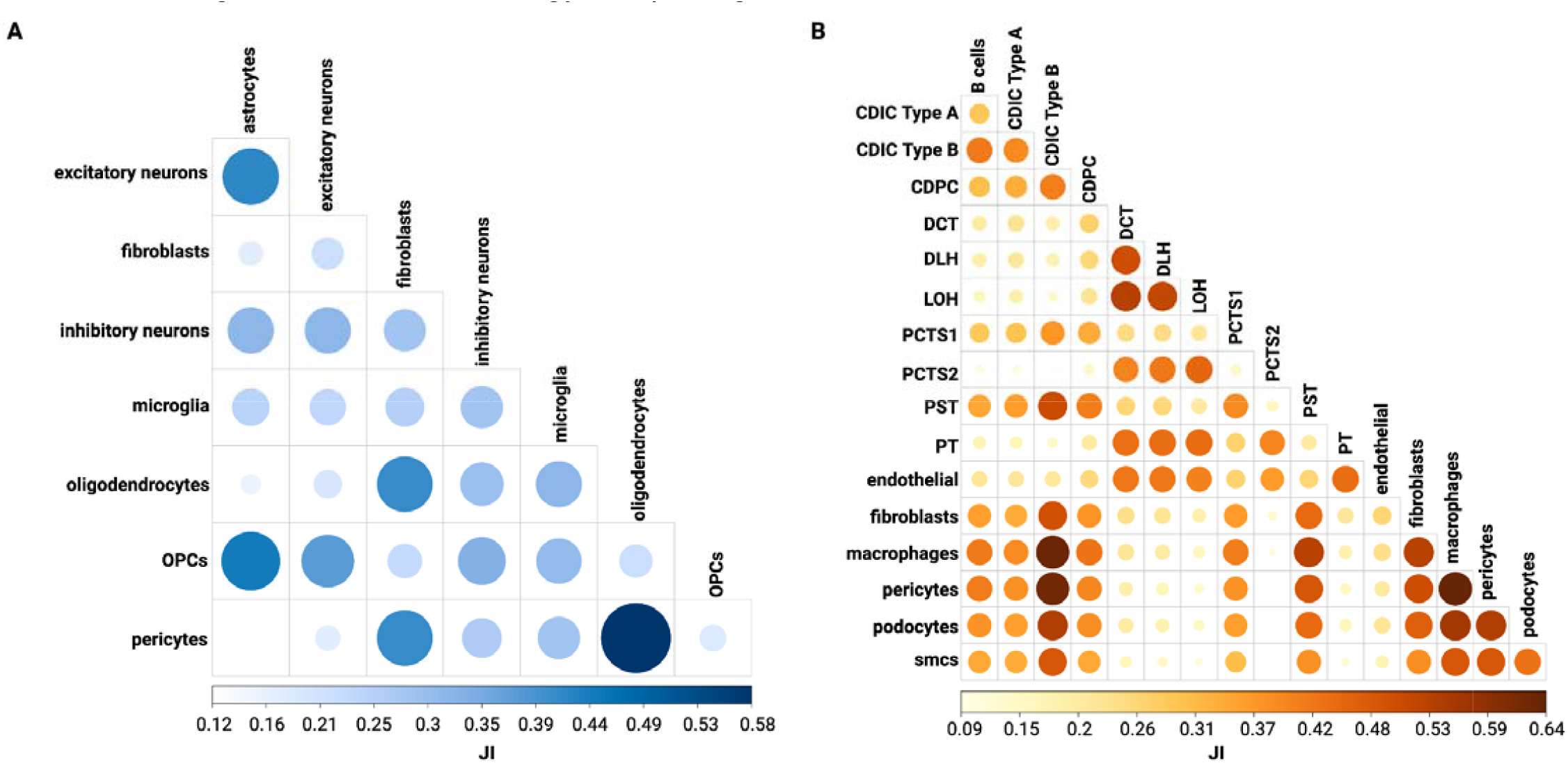
Differential community analysis of SETBP1 communities reveals groups of cell types with more similar SETBP1 differential communities: Jaccard Similarity Index (JI) between all SETBP1 differential communities in **(A)** cerebral cortex and **(B)** kidney.

### Cell-type-specific DNA damage and cell cycle network rewiring in S858R cerebral cortex and kidney cells

While little is known about the role of specific patient variants, several mechanisms are hypothesized to contribute to altered neurodevelopment in SGS, including chromatin remodeling, disrupted cell cycle control, increased DNA damage, and modified PP2A complex activity.^9^ To further evaluate the molecular impact of the S858R variant, we investigated the magnitude and direction of the regulatory network and changes in cooperativity network edge weights where SETBP1 is acting as a TF. Similar to the regulatory network, a more positive edge weight between two proteins indicates a greater likelihood they cooperate to regulate gene expression. In contrast, a more negative edge weight indicates greater confidence that the two proteins do not cooperate. We used these regulatory or cooperativity edge weight magnitude and direction changes to infer potential regulatory rewiring due to the S858R variant. While altered cell cycling, DNA damage, chromatin remodeling, and phosphorylation have been previously implicated by disrupted SETBP1 interactions in the brain^9^, we do not capture SETBP1 interacting with proteins associated with chromatin remodeling and phosphorylation but do identify interactions with proteins related to DNA damage and cell cycle interactions in our networks (**Fig 6A**). We filtered edges to include only those involving SETBP1 as a TF where the target is also a protein in the cooperativity network (these interactions were previously measured by ChIP-seq^5^) and then filtered for APEX1 (cooperating protein from the proposed DNA damage mechanism) and TRP53 (cooperating protein from the proposed cell cycle mechanism), therefore identifying categories depending on the changes in edge weight and direction in the regulatory and cooperativity networks (**Fig 6B**).

**Figure 6:**
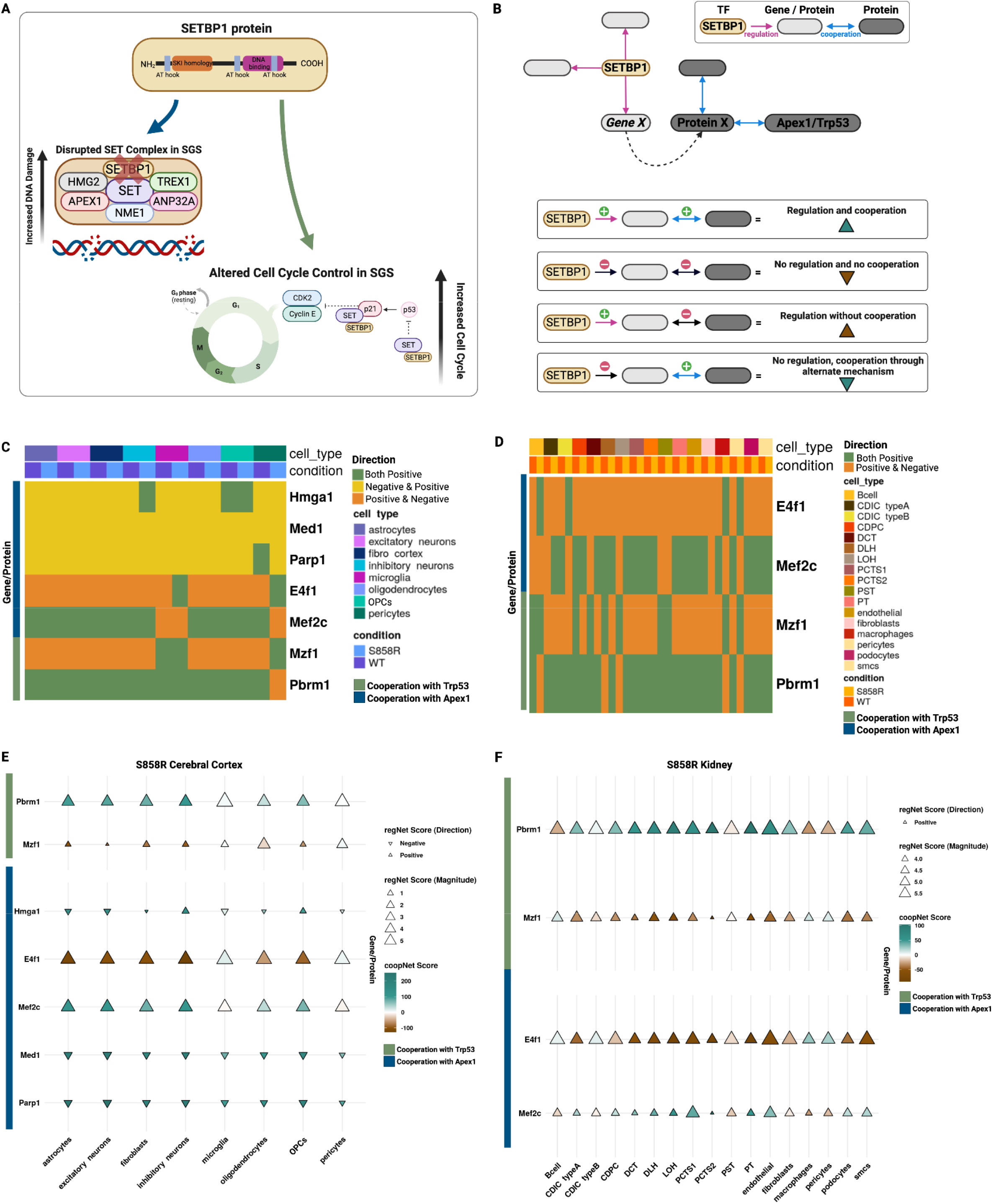
S858R pericytes and microglia and multiple kidney cell types exhibit altered cooperation and regulation in proposed cell cycle and DNA damage mechanisms in SGS: Changes in regulation and cooperation of cell types in S858R cerebral cortex and kidney. **(A)** Schematic of proposed increased DNA damage and increased cell cycle mechanisms in the context of SGS. **(B)** Classification of altered regulation and cooperation through a change in network scores between TF SETBP1 (yellow) and its target genes from the gene set (light grey) associated with proposed mechanisms of altered brain development in SGS mediated through Apex1 or Trp53 (dark grey) for the cerebral cortex. Heatmap showing the sign change in regulation and cooperation between conditions in the cerebral cortex **(C)** and kidney (**D)**. Dot plot showing the Magnitude of regulatory (direction and size of triangle) and cooperativity (teal to brown) network scores in the cerebral cortex **(E)** and kidney **(F)**.

For both the cerebral cortex and kidney, the proposed altered DNA damage and cell cycle mechanisms were cell-type-specific **(Fig 6 C-F, Figure S18)**. We found evidence of several rewiring events. For example, SETBP1 regulation of *Hmga1* is present in S858R but not WT inhibitory neurons, and HMGA1 subsequently cooperates with APEX1, a protein involved in DNA damage. Microglia also had regulatory changes indicative of altered DNA damage mediated through a SETBP1-*E4f1*/E4F1-APEX1 interaction. However, a lack of cooperation between E4F1 and APEX1 is introduced in the S858R condition **(Fig 6C)**. In addition to rewiring involving DNA damage mechanisms mediated through *Parp1*/PARP1, *E4f1*/E4F1, and *Mef2c/MEF2C*, cerebral cortex pericytes also exhibited rewiring for altered cell cycling through *Mxf1*/MXF1 and *Pbrm1*/PBRM1 with *Trp53/TRP53* (the mouse ortholog for human TP53). While the kidney had fewer SETBP1 targets involved in altered mechanisms of cooperation and regulation, more cell types were disrupted. Macrophages, CDIC type B cells, pericytes, B cells, LOH, and DCT all had rewiring involving APEX1 and TRP53 in S858R cells. However, fibroblasts only had regulatory changes involving DNA damage rewiring, represented by a loss of *Mef2c* APEX1 cooperation in S858R compared to WT. CDPC and DLH exhibited SETBP1-*Mzf1/MZF1*-TRP53 mediated cell cycle rewiring through gained cooperation between MZF1 and TRP53 in the S858R compared to WT. Since cooperation does not infer direct interaction or binding, we confirmed interactions for all proteins above using scores from STRING (**Table S8)**.

## Discussion

We profiled the impact of *Setbp1*^S858R^ on cell-type-specific expression and regulation within mouse cerebral cortex and kidney. While previous research primarily focused on the role of NPCs in driving SGS neurodegeneration through aberrant proliferation, deregulated oncogenes and tumor suppressors, unresolved DNA damage, and apoptosis resistance as a result of SETBP1 accumulation in the brain^4,5,9,10^, here we investigated S858R’s impact on multiple cell types in both the cerebral cortex and kidney showing these properties extend to additional cell types. *Setbp1* is known to be ubiquitously expressed across cells and tissues of the body, and we found this to be the case in our study. *Setbp1* was only differentially expressed between *Setbp1*^S858R^ and WT excitatory neurons, but many known targets of SETBP1 were differentially expressed and regulated in multiple cerebral cortex and kidney cell types. Our findings suggest the molecular drivers underlying neurodevelopmental phenotypes in classical SGS also drive atypical SGS, persist after birth, and are present in the kidney. Further, our results, in aggregate, indicate that SETBP1’s role as an epigenetic hub leads to widespread cell-type-specific differences in TF activity, gene targeting, and regulatory rewiring of both cerebral cortex and kidney cells.

Banfi et al. 2021, previously showed that SGS inhibitory neurons inherited DNA damage and were prone to degeneration that could be alleviated by inhibiting PARP-1.^10^ Our rewiring analyses indicated that *Setbp1*^S858R^ pericytes and microglia, in addition to inhibitory neurons, had SETBP1 targets that exhibited altered cooperation with APEX1, potentially identifying additional cell types that may benefit from PARP1 inhibition in SGS.^10^ Cortical pericytes were the only cell type involving TRP53-associated cell cycling mediated through MZF1 and PBRM1. They exhibited the most rewiring across all mechanisms in the cerebral cortex, further emphasizing a potential role for pericytes in atypical SGS. While more cell types involve DNA damage mechanisms mediated through altered cooperation with APEX1 in the kidney, less is known about how this may contribute to a hydronephrosis phenotype. Further functional studies investigating the role of cell-type-specific DNA damage and altered cell cycling would be useful for understanding both the SGS mechanism and treatment opportunities.

There are several limitations to the present study. For example, here we focused on tissues from 6-week-old male mice, so it remains to be seen how cell-type-specific expression and regulation change across developmental time points and in female mice. Furthermore, our TF activity was inferred based on TFs included in the CollecTRI prior. PANDA relied on information from STRING, CIS-BP, and prior SETBP1 ChIP-Seq experiments. As more comprehensive data sets become available, our analytical approaches can be expanded and refined. One major benefit to the TF-gene regulatory networks is that they provide a comprehensive understanding of the impact of a point mutation not just at the gene coexpression or protein-protein cooperation levels but facilitate in silico profiling of *Setbp1*^S858R^ mice across all three facets of the central dogma of biology. However, further experiments are needed to determine if the absence of alterations in SGS-associated phosphorylation and chromatin remodeling mechanisms in regulatory and cooperativity rewiring we found were a direct result of the S858R variant or if this was due to a lack of prior knowledge in the regulatory networks. Additional experiments are also needed to confirm the role of each cell type in SGS etiology and progression. Finally, our study was performed in mice, and while the protein domain is conserved between mouse and human, findings in mouse models do not always translate to patients.

In summary, we report that classical SGS molecular signatures are present across cerebral cortex and kidney cell types in the *Setbp1*^S858R^ mouse model of atypical SGS, underscoring the multifaceted role of SETBP1 and the impact of many cell types in atypical SGS. Future work is needed to further understand how SETBP1’s role changes across cell types, developmental time points, and in the context of other patient variants. Further, experiments and analyses investigating disrupted cell-cell communication between cell types (e.g., astrocytes and excitatory neurons) may reveal novel insights into the role additional cell types play in contributing to SGS neurodegeneration and clinical manifestations outside of the brain.

## Acknowledgments

The authors thank the Lasseigne Lab members Amanda Clark, Anisha Haldar, Avery Williams, Emma Jones, Nathaniel DeVoss, and Victoria Flanary for their feedback throughout this study. Furthermore, they thank Dr. Anna Thalacker-Mercer’s lab at UAB for providing equipment and reagents for ELISA and BCA experiments and Dalton Patterson for his advice for these experiments. In addition, we thank Shanrun Liu at the UAB CFCC (supported by the Center for AIDS Research grant AI027767 and The O’Neal Comprehensive Cancer Center grant CA013148) for his expertise and preparation of snRNA-seq libraries for this study as well as Dr. Michael Crowley and team at the UAB Heflin Center for Genomic Sciences at the UAB Sequencing Core for sequencing the samples. We also thank the UAB Biological Data Science group (RRID:SCR_021766) for providing a script for helping to run containers on the UAB high-performance cluster (https://github.com/U-BDS/training_guides/blob/main/run_rstudio_singularity.sh).

## Authors’ contributions

JHW, VHO, and BNL conceptualized the project. *Setbp1*^S858R^ nuclei isolation and snRNA-seq data were performed and generated by JHW, TMS, and TCH. JHW and TCH performed protein quantification experiments. JHW conducted preliminary analysis and data processing. JHW adapted the original code from TMS and EJW for functional enrichment analysis. VHO aided in downstream conceptualization for analysis of networks and provided initial code for decoupleR transcription factor activity investigation and Jaccard similarity analysis on ALPACA outputs. All other analyses were coded and performed by JHW. TMS, VHO, and TCH reviewed and validated the code for reproducibility. BNL provided supervision, project administration, and funding acquisition. JHW wrote the first draft. JHW, TMS, TCH, EJW, VHO, and BNL reviewed and edited manuscript versions. All authors read and approved the final manuscript.

## Appendices

Whitlock_SUPP_figures&tables_etal_CellularMolecularMedicine_2023.docx

Whitlock_SUPP_Methods_etal_CellularMolecularMedicine_2023.docx

## Supporting information

Table_S3.xlsx

Table_S4.xlsx

Table_S5.xlsx

Table_S6.xlsx

Table_S7.xlsx

## Notes

**Data and code availability statement:** Data and code supporting the results of this study are available at Zenodo (doi: 10.5281/zenodo.8192482) and https://github.com/lasseignelab/230227_JW_Setbp1Manuscript (doi:10.5281/zenodo.8190948). Docker images used for these analysis are publicly available on Docker Hub and Zenodo (doi:10.5281/zenodo.8190923).

**Funding statement:** This work was supported in part by the UAB Lasseigne Lab funds (to BNL), UAB Pilot Center for Precision Animal Modeling (C-PAM)(1U54OD030167) (to BNL), JHW was funded by the UAB Predoctoral Training Grant in Cell, Molecular, and Developmental Biology (CMDB T32)(5T32GM008111-35), TMS was funded by the Alzheimer’s of Central Alabama Lindy Harrell Predoctoral Scholar Program.

**Conflict of interest disclosure:** The authors have declared no conflicts of interest

https://doi.org/10.5281/zenodo.8190948

https://doi.org/10.5281/zenodo.8190923

https://doi.org/10.5281/zenodo.8192482

### Competing Interest Statement

The authors have declared no competing interest.

