## Supplemental Methods for "Cell-type-specific gene expression and regulation in the cerebral cortex and kidney of atypical *Setbp1*^S858R^ Schinzel Giedion Syndrome mice"

### Supplementary Materials and Methods:

##### ***In-vivo* Mouse Samples**

Jackson Laboratories (JAX) snap-froze tissues with liquid nitrogen upon collection, and we stored these samples at -70ºC upon receipt until nuclei isolation.

##### **Kidney Nuclei Isolation**

We performed single nuclei isolation for the kidney and adapted our protocol from the one previously described by Kirita et al. *PNAS* 2020.^1^ We isolated nuclei using the Nuclei EZ Prep kit with Nuclei EZ Lysis buffer (Sigma #NUC-101) supplemented with cOmplete ULTRA tablets (Sigma# 05892791001), SUPERase IN (Thermosisher# AM2696), and Promega RNasin Plus (Fisher Promega #PRN2615). We minced whole kidney cortex into <1mm pieces and then homogenized them using a Dounce tissue grinder (Kimble# 8853000002) in 2 ml of cold Nuclei EZ Lysis buffer. We then filtered the homogenate through a 200-µm strainer (Pluriselect#:43-50200) and a 40-µm strainer (Pluriselect#:43-50040) and then centrifuged at 500 x g for 5 min at 4 ºC. We resuspended the pellet in 4ml of the buffer and incubated it on ice for 5 minutes. After centrifugation at the same time, temperature, and speed, we resuspended the pellet in Nuclei Suspension Buffer as described by Kirita et al. *PNAS* 2020.^1^ We counted nuclei on the C-Chip (Fisher# 22600103) to obtain a final count of 1.3 - 3.0 million nuclei captured per kidney.

##### **snRNA-seq processing**

Per the standard 10X protocol, the UAB CFCC used the 10X Chromium Controller to partition kidney and cerebral cortex nuclei into droplets with a barcoded gel bead where nuclei were lysed, and RNA was reverse transcribed, creating cDNA encapsulated in each droplet. After fragmenting and amplifying cDNA, they added Illumina adapters using the Chromium Single Cell 3ʹ GEM, Library & Gel Bead Kit v3 (10X#: PN-1000121). The UAB Heflin Center for Genomic Sciences at the UAB Sequencing Core indexed and sequenced all samples on a NovaSeq 6000 (Illumina). For each sample, they loaded approximately 20,000 nuclei onto an S4 flow cell.

###### **Data pre-processing**

We processed the snRNA-seq data with 10X Genomics Cell Ranger (version 6.1.1) according to 10X Genomics standard protocols. We aligned sequencing reads to *mus musculus* (mm10) to generate feature-barcode matrices. The average paired-end sequencing depth was 81,000 reads per nuclei, with approximately 12,000 and 8,000 nuclei per sample for the kidney and cerebral cortex, respectively. We investigated raw sequencing data quality using FastQC (version 0.11.9).^2^

###### **Variant Confirmation**

Prior to receipt of samples, JAX confirmed the Setbp1 S858R variant using qPCR to ensure the presence of C/G, representing the coding change in heterozygotes, compared to C/C in WT. We additionally confirmed all variants post-sequencing. All heterozygous samples had the S858R variant present **(Figure S1)**.^3^

###### **Setbp1 Protein Quantification**

To quantify SETBP1 protein abundance in kidney and cerebral cortex samples, we combined approximately 20 mg of input tissue with a 1:10 dilution of 1X PBS and homogenized in both tissues using the same homogenization protocol we used for cortex nuclei isolation. Following the MOUSE SETBP1 ELISA Kit (MyBioSource Cat No. MBS9335445), we centrifuged the homogenates at 1000 x g (3000rpm) for 20 minutes at 4ºC. We then loaded supernatant and appropriate blanks into a pre-coated ELISA microwell plate. We performed incubation and manual washing as described in the MyBioSource Kit protocol and measured absorbance using a Tecan Infinite M Plex plate reader at 450 nm for 10 flashes. By fitting the absorbance with a standard linear regression, we quantified SETBP1 concentration in սg/mL at 450 nm. In addition, we performed a BCA on all samples following the Pierce protocol.^4^ (**Figure S2**)

**Single-nuclei RNA-sequencing processing:**

All downstream data processing was done using R version 4.1.3 unless otherwise specified.

###### **Data processing and quality control**

We read the gene barcode matrices with the Read10X function in Seurat (version 4.3.0). We annotated metadata, including the condition and sample ID for cell barcodes. Next, we constructed Seurat objects using the CreateSeuratObject function for each sample from the raw gene-barcode matrices. Then, we merged all objects for each dataset using the MergeSeurat function. We removed all mitochondrial genes transcribed from the mitochondrial genome in all datasets consistent with snRNA-Seq best practices.^5,6^ Due to the high sequencing depth, we filtered for nuclei in either tissue with mitochondrial reads <5% and where nFeature_RNA was >=1000 and <= 15000. We did not filter nCount_RNA. A large proportion of reads in all samples mapped to *Malat1*, a known marker of extranuclear RNA contamination, so we removed these reads.^7^ We examined the cell cycle stage and saw that the nuclei appeared evenly distributed between G2M and S phases across tissues and conditions; therefore, we did not regress out cell cycle signatures.

We converted Seurat objects to log scale (Seurat function NormalizeData) and scaled the data, regressing out percent_mito and nFeature_RNA (Seurat function ScaleData). We used FindVariableGenes from Seurat to exclude lowly expressed genes from the data set below a nFeature of 4,000.

###### **Clustering, dimension reduction, and cell type identification**

###### Using Seurat’s ElbowPlot function, we selected 30 PCs for each dataset and integrated them using Harmony (version 0.1.1).^8^^,^^9^ We clustered the Seurat objects from each dataset using the Seurat FindNeighbors, and FindClusters commands from resolutions 0.5 - 2.2. by Leiden clustering.^10^ We visualized relationships between clusters across resolutions using clustree (version 0.5.0) and selected the most stable resolutions.^11^ We clustered the cerebral cortex and kidney data at a resolution of 1.0 and 1.7, respectively, for use in all downstream analyses. We detected differentially expressed genes (DEGs) between clusters using the Seurat FindAllMarkers function with a Wilcoxon test on the 'RNA' assay, specifying only genes with a log2FC cutoff greater than or equal to 0.2 for both datasets. We compared top marker genes for each cluster against markers identified from the literature and PanglaoDB, a community-curated cell-type marker database, to assign cell types (**Table S2)**.^12–17^

###### **Ambient RNA removal**

Droplet-based sc/snRNA-seq is susceptible to cell-free RNA that can confound the biological interpretation of transcriptome data.^18^ This extranuclear or ambient RNA is even more prevalent in complex samples, such as kidney tissue; because of this, we used SoupX^19^ (version 1.6.2) to determine the fraction of ambient RNA in each single nuclei. After estimating contamination, we removed the contaminated nuclei. We then re-ran the Seurat pipeline as previously described. After re-clustering, the final resolution for kidney samples was 1.5.

###### **SETBP1 target gene set construction**

We constructed a gene set of *Setbp1* and its known targets using experimental ChIP-seq binding sites of SETBP1 provided by GTRD^20^ ([GSM2300423](https://nam12.safelinks.protection.outlook.com/?url=http%3A%2F%2Fwww.ncbi.nlm.nih.gov%2Fgeo%2Fquery%2Facc.cgi%3Facc%3DGSM2300423&data=05%7C01%7Cjwhitlock%40uab.edu%7Ced4e34457a9c4581007908db0496423e%7Cd8999fe476af40b3b4351d8977abc08c%7C1%7C1%7C638108815651478132%7CUnknown%7CTWFpbGZsb3d8eyJWIjoiMC4wLjAwMDAiLCJQIjoiV2luMzIiLCJBTiI6Ik1haWwiLCJXVCI6Mn0%3D%7C3000%7C%7C%7C&sdata=59LpWrO6D0VEXMujxXgDNnJ8iPF9jggjY1mFAPHTGV4%3D&reserved=0), [GSE86334](https://nam12.safelinks.protection.outlook.com/?url=http%3A%2F%2Fwww.ncbi.nlm.nih.gov%2Fgeo%2Fquery%2Facc.cgi%3Facc%3DGSE86334&data=05%7C01%7Cjwhitlock%40uab.edu%7Ced4e34457a9c4581007908db0496423e%7Cd8999fe476af40b3b4351d8977abc08c%7C1%7C1%7C638108815651478132%7CUnknown%7CTWFpbGZsb3d8eyJWIjoiMC4wLjAwMDAiLCJQIjoiV2luMzIiLCJBTiI6Ik1haWwiLCJXVCI6Mn0%3D%7C3000%7C%7C%7C&sdata=Ob3Nsm6tNPlYXjW7PInGSL%2Bovk8bK5HtYNHW8auDGYU%3D&reserved=0)), curated databases such as MSigDB^21^ (M40860), SIGNOR^22^ (version 3.0; accessed January 2023), TRRUST^23^ (version 2.0; accessed January 2023) and literature mining^24,25^. We identified a total of 226 human genes mapped to 682 mouse orthologs using bioDBnet^26^ (version 2.1) **(Table S5).**

###### **Investigation of the presence of NPCs, reactive astrocyte subclusters, and expression of *Set*, *Ppp2ca*, and failed repair-associated genes**

We extracted astrocytes based on active.ident cell identity metadata in the annotated Seurat object and re-clustered using the same clustering methods above at a resolution of 0.9. We determined reactive astrocyte presence in subclusters using UMAP dimensional reduction paired with FeaturePlot expression of *Gfap* (a known marker of reactive astrocytes in Alzheimer's disease (AD) and other neurodegenerative diseases^27^) between conditions **(Figure S14)**. We used Seurat’s FindAllMarkers function to identify differentially expressed marker genes between astrocyte subclusters using the same methods above for processing the entire single-nuclei object. In addition, we also generated violin plots of proliferating NPC marker, *Sox2*, examined the expression of *Ppp2ca* and *Set*, known to interact with SETBP1, and expression of kidney genes associated with failed repair (*Vcam1*, *Havcr1*, *Krt20*, *Myc*, *Slc22a30*, *Sox9,* *Dcdc2a*, *Slc5a12,* and *Slc7a13*) **(Figures 2E, S12, S15, and S16)**.^1,28,29^

###### **TF activity**

Using decoupleR^30^ (version 2.6.0), we inferred TF activity **(Figure 3)**. In our analysis, we combined scRNA-seq data with prior knowledge. To map prior biological knowledge, we used pseudo-bulked gene by sample matrices for each cell type as input of molecular readouts and CollecTRI, a comprehensive network of TFs and their direction of regulation on transcriptional targets (accessed May 2023).^31^ We calculated activity scores for all TFs that had a minimum of 5 targets using the Multivariate Linear Model (run_mlm), where the t-values from the model represent regulator activity.^30^ We scaled and centered data using Seurat ScaleData and calculated average scores for every TF. To prioritize TFs with the largest change in activity between conditions, we calculated the absolute percentage change of the average activity for every TF between conditions. We thresholded absolute percent changes to only include those above the third quartile or below the first quartile for both tissues.

##### **Aggregate Network Analysis**

We constructed cell-type-specific networks using Passing Attributes between Networks for Data Assimilation (PANDA) framework from the netZooR package (version 1.2.1) to investigate the cell-type-specific impact of the S858R variant on regulation.^32,33^ PANDA requires multi-omic inputs in the form of gene expression, protein-protein interaction (PPI), and TF-motif binding data. We constructed PANDA networks using a docker container (**see availability statement**) running R version 4.2.1 in the intersection mode, meaning any nonintersecting gene and TF sets were not included in the resulting network. We included information on average run time, number of iterations to build each network, number of genes and TFs measured, and final hamming distance for each cell-type-specific regulatory network in **Table S6**.

###### **Prior coexpression network based on self-generated expression data**

###### **Pseudobulk snRNA-seq**

To overcome data sparsity, we pseudo-bulked the snRNA-Seq expression data and generated count matrices for each cell type per sample. To achieve this, we extracted sample-specific count matrices from the Seurat objects using the normalized 'data' from the 'RNA' slot for every gene and cell and then aggregated per gene for each cell type across samples into a new matrix, transforming from cell x gene counts per sample to gene x sample counts for each cell type.^6^ We used the newly transformed matrices as an input for the cell-type-specific network generation described below.

###### **Prior regulatory network based on TF Motif Information**

We obtained TF-motif binding input data from Sonawane et al. Cell Reports, 2017, which was previously constructed using the Catalog of Inferred Sequence Binding Preferences (CIS-BP).^34,35^ We used it to generate a prior regulatory network with PANDA.^34,35^^,^^32^ This *mus musculus* motif prior included direct and inferred evidence for associated DNA binding motifs. We mapped motif IDs from CIS-BP to motifs in the RefSeq prior regulatory network (accessed October 1, 2022; <https://sites.google.com/a/channing.harvard.edu/kimberlyglass/tools/resources>) containing the -750 and +750 nucleotides from all TSS in the mm10 genome. We enriched the TF motif resource with the SETBP1 target gene set. This process resulted in an initial map of potential TF-motif regulatory interactions involving 1294 unique TFs targeting 24,556 unique motifs in mouse. We excluded three genes (ENSMUSG00000090020, ENSMUSG00000079994, and ENSMUSG00000044690) in motif mappings as they did not have an associated gene symbol.

###### **Prior Protein-Protein Interaction Network**

We estimated the PPI network using interaction scores for *mus musculus* (species.index: 10090) in STRING (version 11.0) (accessed with STRINGdb R package version 2.6.5 on August 5, 2022).^36^ STRING PPI input data has different score thresholds, corresponding to varying confidence levels in STRING interactions based on experimental data, computational prediction methods, and public text collections. We used all of STRING to ensure we were not excluding large numbers of interactions and provided the opportunity to capture novel and understudied interactions. We did not include missing interactions (TF motif and gene interaction = 0). This resulted in 338,752 interactions between 1,118 unique proteins for mice (less than 1% of identifiers did not map).

###### **Gene Targeting Score Calculation**

We filtered all regulatory networks for the cerebral cortex (n = 16; 8 cell types across two conditions) and kidney cortex (n = 34; 17 cell types across two conditions) for only positive edge weights and calculated targeting scores for all genes within cell types for S858R and WT by summing the edge weights of all inbound regulatory edges for a gene based on the indegree.^37^

###### **Cell-type-specific Differential Gene Targeting**

We calculated differential targeting between conditions by taking the difference in targeting scores for each gene between S858R and WT conditions, where the more positive targeting scores represent genes enriched in the S858R mice. The closer a value is to 0, the less difference in targeting between conditions for that cell type. We performed differential targeting on all genes, split scores into positive and negative, and selected everything above the third quartile in each direction for each cell type. Once scores were thresholded, we investigated the global landscape of gene targeting by summing all targeting scores by cell type, normalized by the total number of genes. We ranked genes by differential targeting score and subset for the top 20 within each cell type. Differentially targeted genes passing the above quantile thresholds were filtered for those within the SETBP1 gene set **(Figure 4A, 4B)**. In addition, we performed FEA using GO and HP enrichment of differentially targeted genes in the SETBP1 gene set, showing increased targeting in the S858R condition after setting a p-adjusted threshold of 0.05 using a Bonferroni procedure for multiple hypothesis correction **(Figure 4C, 4D)**.

###### **Investigation of proposed mechanisms of altered neurodevelopment in SGS through cooperativity**

The PANDA algorithm provides three network outputs: a regulatory network modeling TF-gene interactions, a cooperativity network modeling protein-protein cooperation, and a coexpression network of gene-gene relationships. From these, we filtered all regulatory networks for TF, SETBP1, and all cooperativity networks for only nodes within the SETBP1 target gene set. We mapped the nodes in the regulatory network to their corresponding proteins in the cooperativity network to identify all edges where SETBP1 regulates a target gene and that target gene, as a protein, cooperates with another protein in the cooperativity network. We subsetted all proteins in the cooperativity network edges that were not part of the SETBP1 gene set for proteins in previously proposed mechanisms contributing to altered neurodevelopment in Schinzel Giedion Syndrome **(Figure 6)**.^25^

We used the following to annotate the mechanisms:

Altered Cell Cycle: Cdk2, Trp53, Tp53, Tceal1 Set; Chromatin remodeling: Anp32a, Taf1a, Set; DNA damage: Apex1, Anp32a, Apex, Hmg2, Hmgb2, Nme1, Set, Trex1; Phosphorylation: Ppp2r1a, Anp32a, Set, Akt, Akt1.

We annotated the magnitude, either positive or negative, of each edge weight in the regulatory and cooperativity networks for these edges and the sign change between the regulatory network and the cooperativity network edges as one of the following: positive-positive (regulation and cooperation occurring); positive - negative (regulation occurring and no cooperation); negative - negative no regulation and no cooperation); and negative - positive (no regulation, and cooperation occurring through an alternate mechanism).

1.	Kirita, Y., Wu, H., Uchimura, K., Wilson, P. C. & Humphreys, B. D. Cell profiling of mouse acute kidney injury reveals conserved cellular responses to injury. *bioRxiv* 2020.03.22.002261 (2020) doi:[10.1101/2020.03.22.002261](http://dx.doi.org/10.1101/2020.03.22.002261).

2.	Babraham Bioinformatics - FastQC A Quality Control tool for High Throughput Sequence Data. <https://www.bioinformatics.babraham.ac.uk/projects/fastqc/>.

3.	Robinson, J. T. *et al.* Integrative genomics viewer. *Nat. Biotechnol.* **29**, 24–26 (2011).

4.	MAN0011430_Pierce_BCA_Protein_Asy_UG.pdf.
