## Supplemental Figures and Tables for "Cell-type-specific gene expression and regulation in the cerebral cortex and kidney of atypical *Setbp1*^S858R^ Schinzel Giedion Syndrome mice"

**Table S1: Sample Sheet**

| **sample_ID** | **tissue** | **model/condition** | **age (weeks)** | **sex** |
| --- | --- | --- | --- | --- |
| J1 | right cerebral cortex | S858R | 6 | M |
| J2 | right cerebral cortex | C57BL6/J control | 6 | M |
| J3 | right cerebral cortex | C57BL6/J control | 6 | M |
| J4 | right cerebral cortex | C57BL6/J control | 6 | M |
| J13 | right cerebral cortex | S858R | 6 | M |
| J15 | right cerebral cortex | S858R | 6 | M |
| K1 | right kidney | C57BL6/J control | 6 | M |
| K2 | right kidney | S858R | 6 | M |
| K3 | right kidney | C57BL6/J control | 6 | M |
| K4 | right kidney | S858R | 6 | M |
| K5 | right kidney | C57BL6/J control | 6 | M |
| K6 | right kidney | S858R | 6 | M |

**Table S1:** Sample sheet containing ID, tissue, model, condition, age in weeks, and sex of mice used in this study. Heterozygous, S858R mice were smaller and lighter than *Setbp1* (+/+), with reduced brain, liver, and kidney organ weight.

**Figure S1: S858R Variant confirmation in IGV**

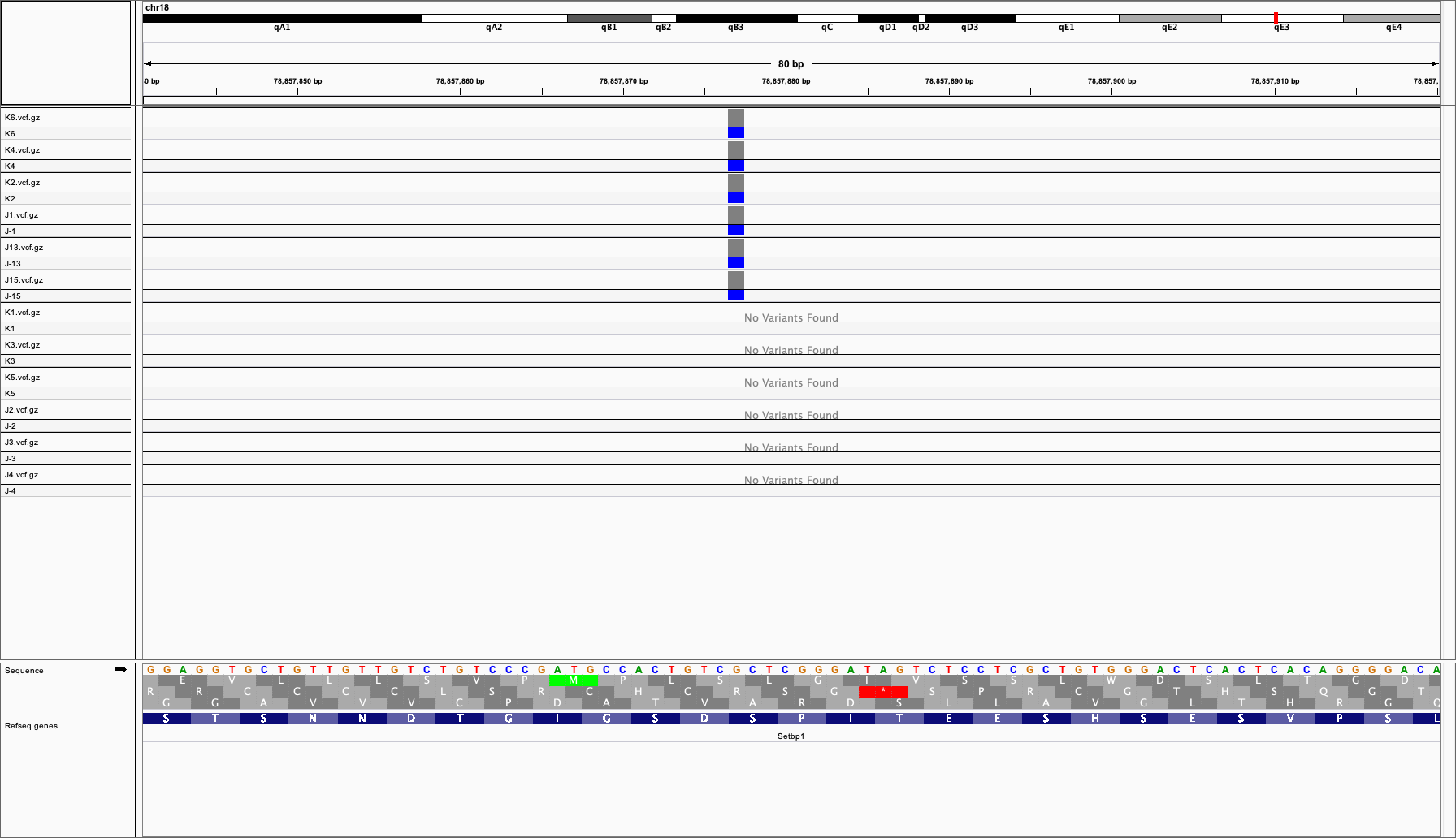

**Figure S1:** Confirmation of variant presence (indicated by blue) using IGV in all heterozygous samples at chr18:78857841-78857920. Samples K1, K3, K5, J2, J3, and J4 are Ctrl for the kidney and brain, respectively. Samples K2, K4, K6, J1, J13, and J15 are heterozygous for S858R in *Setbp1*.

**Figure S2: Setbp1 total protein is significantly increased in S858R compared to WT**

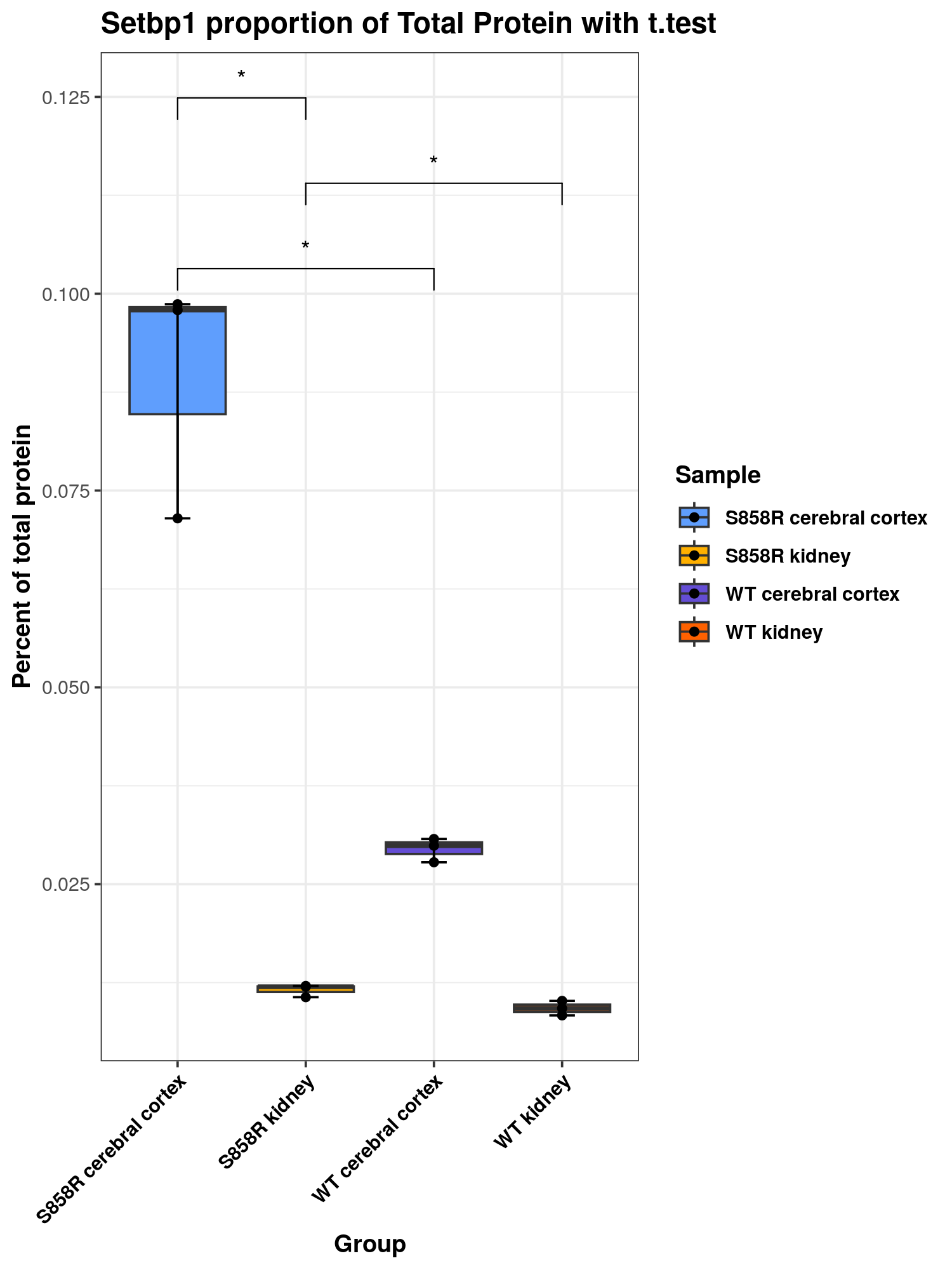

**Figure S2:** Boxplot of the proportion of total protein as a percent (y-axis) in S858R (light blue) cerebral cortex and S858R kidney (yellow) compared to WT cerebral cortex (dark blue) and kidney (orange). Significance was calculated using a paired t.test, and comparison groups with a significant difference in Setbp1 abundance are indicated with an asterisk.

**Table S2: Table of marker genes for cortex and kidney cell types used in cell type annotation**

| **Cerebral Cortex Cell Type** | **Marker Gene(s)** |
| --- | --- |
| Excitatory neurons | *Slc17a7, Pcp4* |
| Inhibitory neurons | *Gad1, Synpr* |
| Oligodendrocytes | *Hapln2, Opalin, Ptgds* |
| Oligodendrocyte precursor cells (Opcs) | *Pdgfra, Olig2* |
| Astrocytes | *Slc1a3, Gja1, Aqp4, Phka1* |
| Microglia | *Cx3cr1, Csf1r, Dock8* |
| Pericytes | *Vtn* |
| Fibroblasts | *Nr4a2, Bnc2* |
| **Kidney Cell Type** | **Marker Gene(s)** |
| Pericytes | *Pdgfrb* |
| Endothelial | *Kdr, Ptprb* |
| Proximal Tubule (PT) | *Slc34a1*, *Slc13a3* |
| Proximal Straight Tubule (PST) | *Slc22a7*, *Atp11a* |
| Proximal Convoluted Tubule Segment 1 (PCTS1) | *Slc5a2, Slc5a12* |
| Proximal Convoluted Tubule Segment 2 (PCTS2) | *Fxyd2* |
| Loop of Henle (LOH) | *Slc12a1* |
| Collecting Duct Principal cells (CDPC) | *Aqp2, Hsd11b2* |
| Distal Convoluted Tubule (DCT) | *Slc12a3* |
| Macrophages | *Runx1, Ptprc* |
| Distal Loop of Henle (DLH) | *Bst1, Akr1b3* |
| Collecting Duct Intercalated cells (CDIC) | *Atp6v1g3, Atp6v0d2* |
| CDIC type A | *Aqp6* |
| CDIC type B | *Hmx2* |
| B cells | *Cd79b, Bank1* |
| Podocytes | *Nphs1, Nphs2, Wt1* |
| Fibroblasts | *Pdgfra* |
| Smooth muscle cells | *Atp1a2* |

**Table S2:** Table containing the marker genes used in this study to annotate cell types for both tissues.

**Tables S3, S4: DEGs for S858R mice in each cell type**

Included in file Table_S3.xlsx and Table_S4.xlsx

**Table S3:** Table containing the differentially expressed genes (DEGs) identified in this study for kidney cell types. Information provided in the table contains each DEG, the p-value, its average log2FC, the percent of nuclei expressing that gene in each condition (pct.1 and pct.2), the adjusted p-value, which condition (cluster) that gene was differentially expressed within, and the cell type.

**Table S4:** Table containing the differentially expressed genes (DEGs) identified in this study for cerebral cortex cell types. Information provided in the table contains each DEG, the p-value, its average log2FC, the percent of nuclei expressing that gene in each condition (pct.1 and pct.2), the adjusted p-value, which condition (cluster) that gene was differentially expressed within, and the cell type.

**Table S5: Table of SETBP1 target gene set**

Included in file Table_S5.xlsx

Table of known targets of SETBP1 from the SETBP1 target gene set we constructed (**See Supplemental Methods)**

**Table S6: PANDA network construction information and run time**

Included in file Table_S6.xlsx

Table containing additional information on PANDA TF-gene regulatory network construction for each cell type in both tissues including the time (seconds), number of iterations needed to reach consensus, total number of genes in the network, number of TFs in the network, and final hamming distance.

**Figure S3: Marker genes for cortex and kidney cell type annotation and proportion of nuclei**

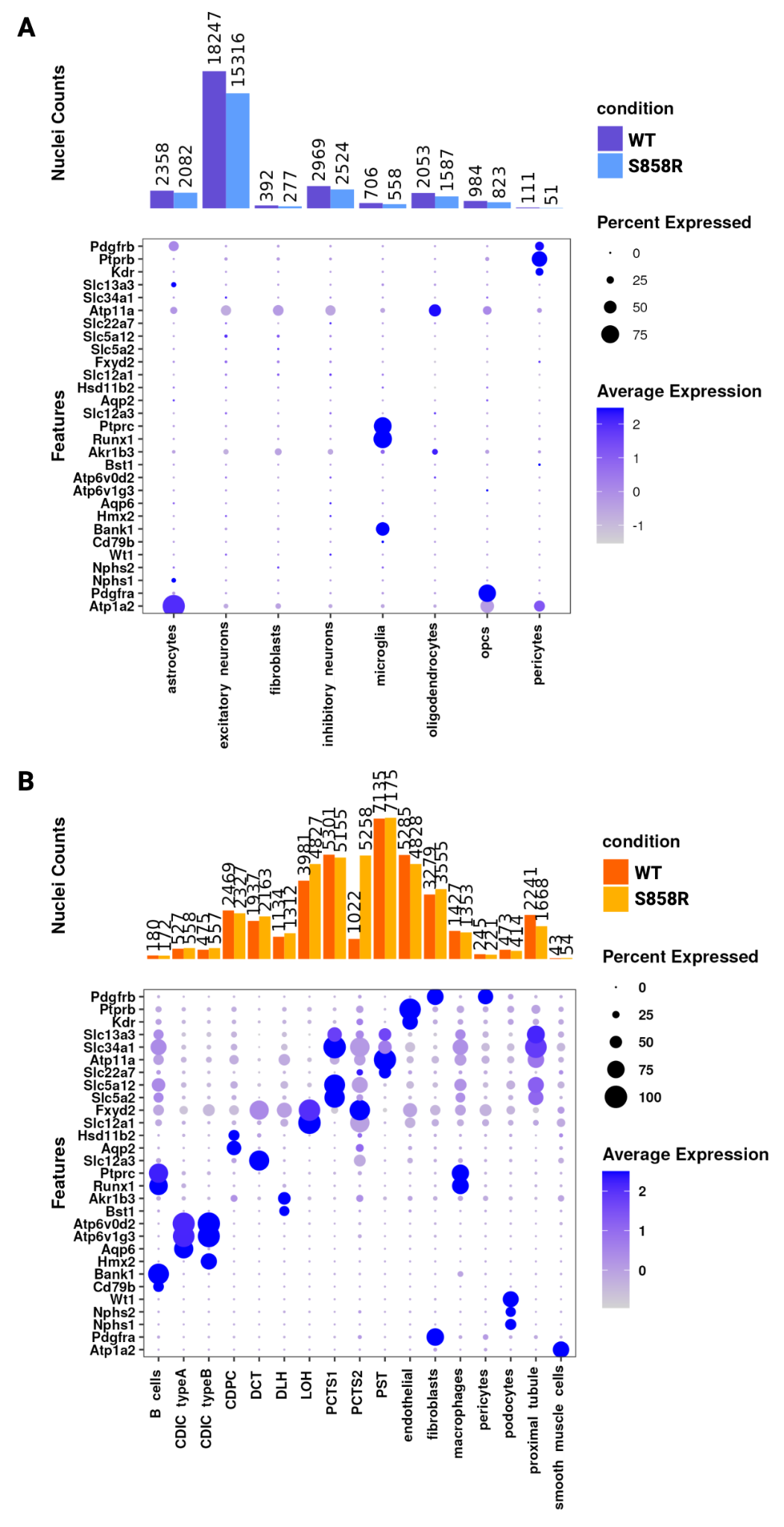

**Figure S3:** Stacked barplot depicting the nuclei counts (y-axis) for each cell type across conditions (top) and dot plot of marker genes used for cell type identification (bottom) for cerebral cortex **(A)** and kidney **(B).** The size of the dot indicates the percent of nuclei expressing that marker and the average expression is represented as a scale from white to dark blue.

**Figure S4: UMAP of Setbp1 S858R snRNA-seq by condition and sample**

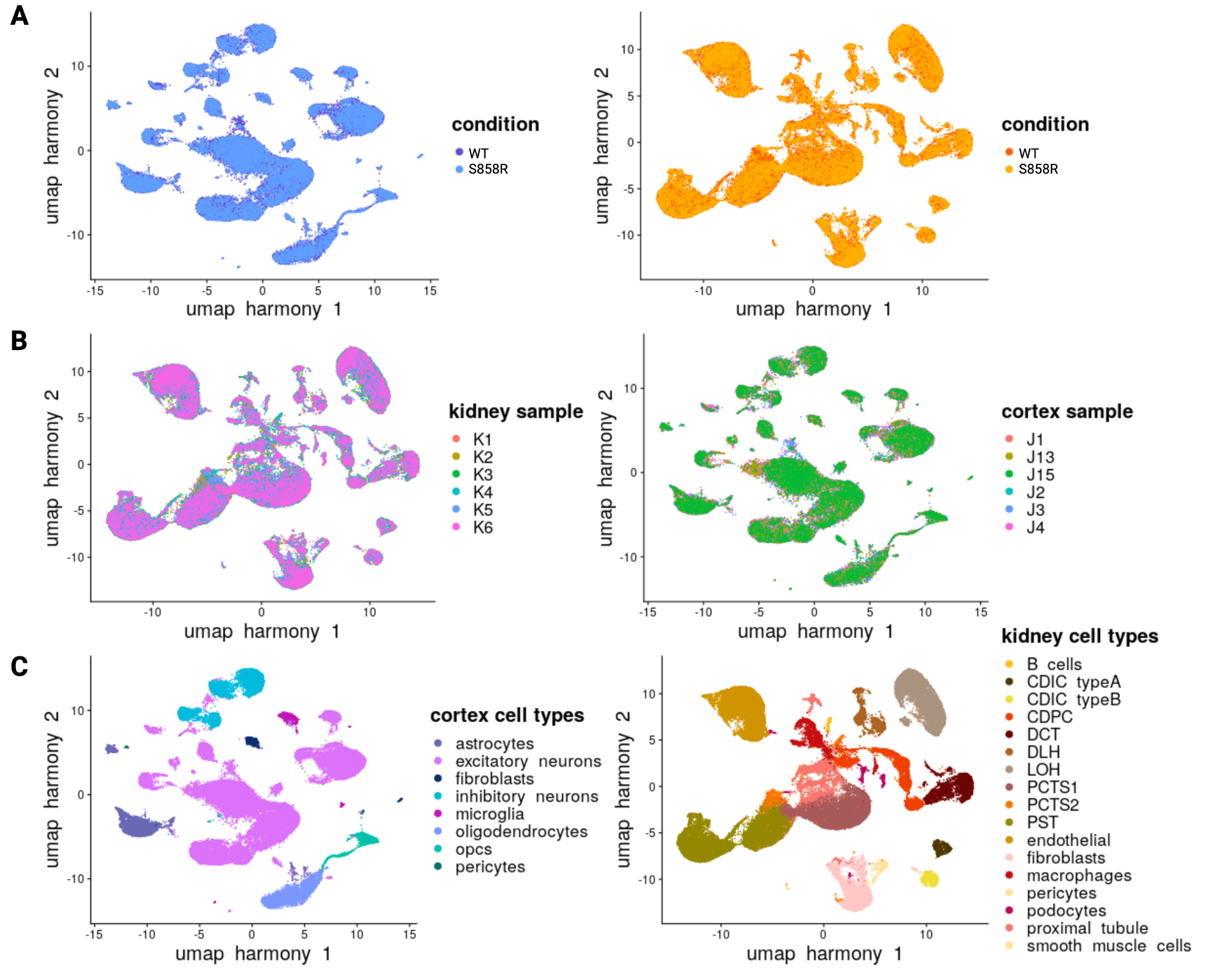

**Figure S4:** Visualization of UMAP dimensional reduction on post-harmony integration snRNA-seq data for both tissues by condition, sample, and cell type.

**Figure S5: gprofiler2 GO pathway analysis on predicted Setbp1 target genes upregulated in S858R excitatory neurons**
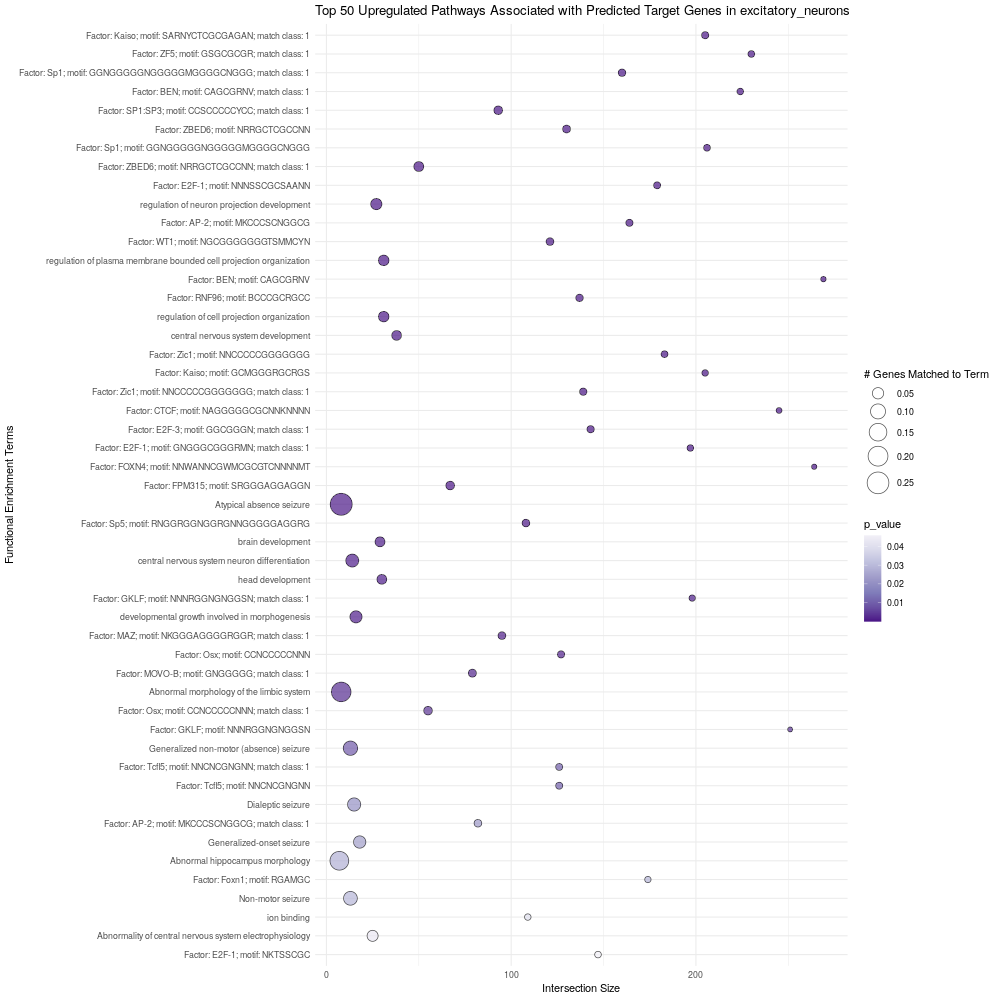

**Figure S5:** Dot plot of the top 50 upregulated pathways (y-axis) associated with SETBP1 target gene set in excitatory neurons. The color of the dot corresponds to the significance (p-value; light purple to dark purple) and the size of the dot to the intersection size (proportion of genes matched to that term).

**Figure S6: gprofiler2 GO pathway analysis on predicted Setbp1 target genes downregulated in S858R excitatory neurons**
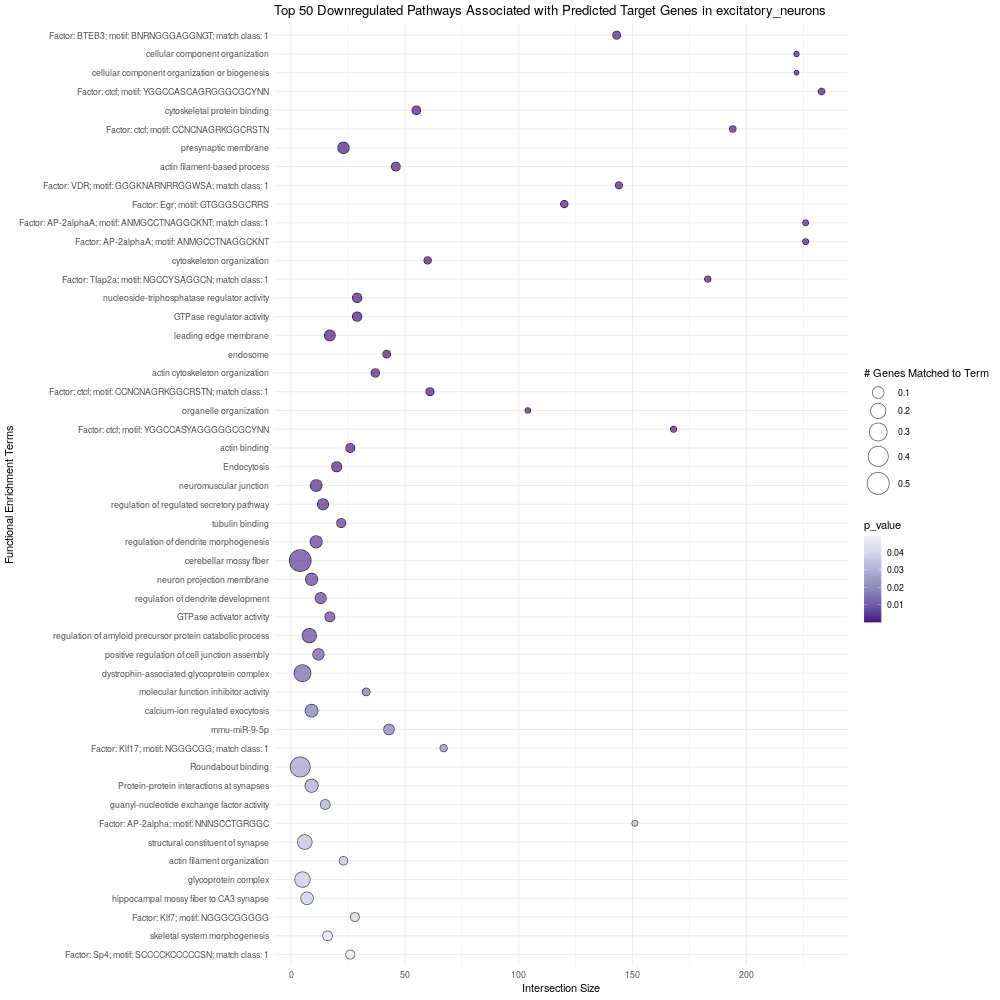

**Figure S6:** Dot plot of the top 50 downregulated pathways (y-axis) associated with SETBP1 target gene set in excitatory neurons. The color of the dot corresponds to the significance (p-value; light purple to dark purple) and the size of the dot to the intersection size (proportion of genes matched to that term).

**Figure S7: gprofiler2 GO pathway analysis on predicted Setbp1 target genes downregulated in S858R astrocytes**

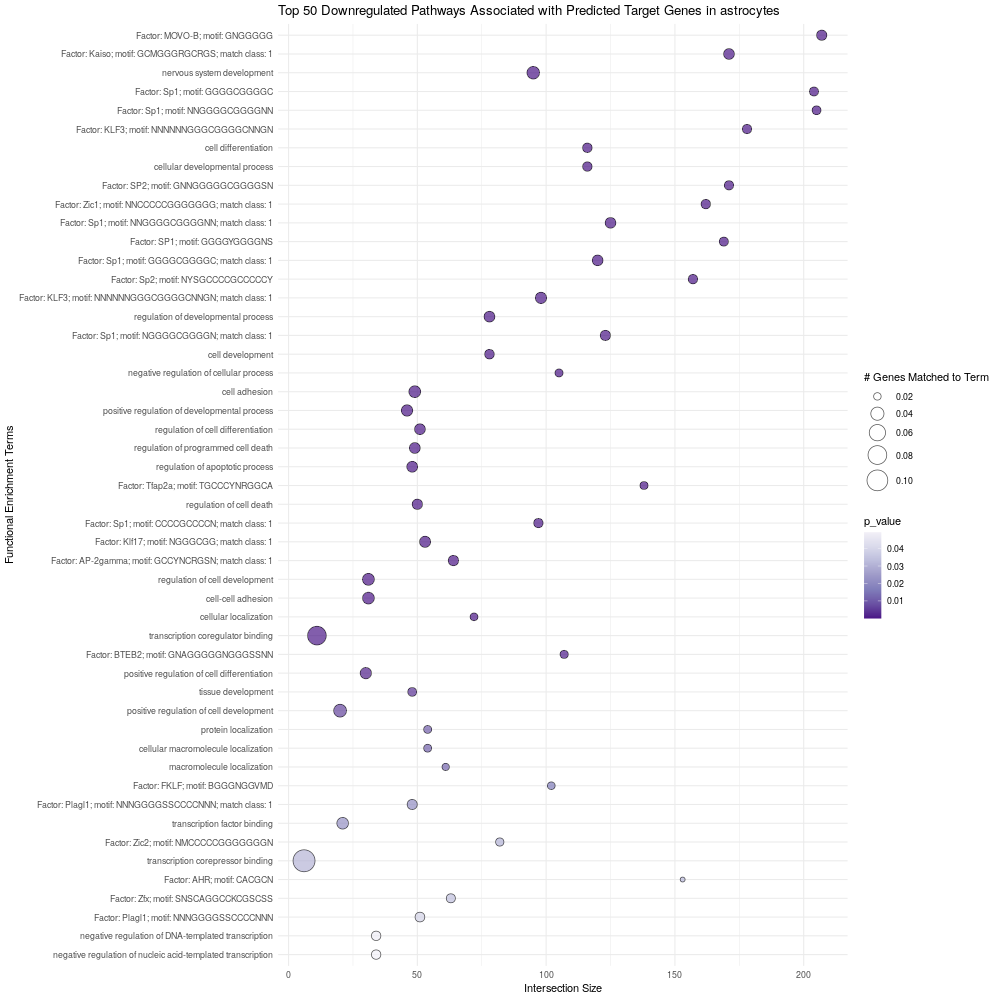

**Figure S7:** Dot plot of the top 50 downregulated pathways (y-axis) associated with SETBP1 target gene set in astrocytes. The color of the dot corresponds to the significance (p-value; light purple to dark purple) and the size of the dot to the intersection size (proportion of genes matched to that term).

**Figure S8: gprofiler2 GO pathway analysis on predicted Setbp1 target genes downregulated in S858R inhibitory neurons**

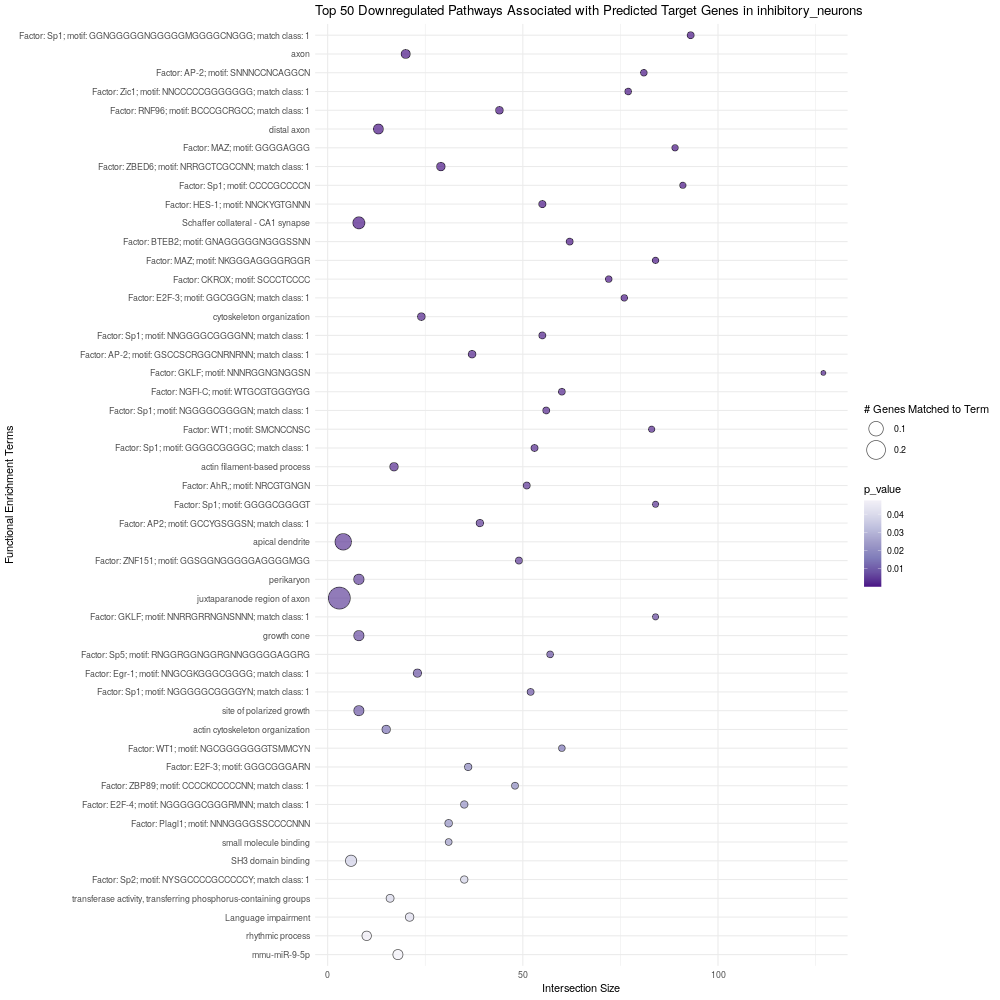

**Figure S8:** Dot plot of the top 50 downregulated pathways (y-axis) associated with SETBP1 target gene set in inhibitory neurons. The color of the dot corresponds to the significance (p-value; light purple to dark purple) and the size of the dot to the intersection size (proportion of genes matched to that term).

**Figure S9: gprofiler2 GO pathway analysis on predicted Setbp1 target genes downregulated in S858R microglia**

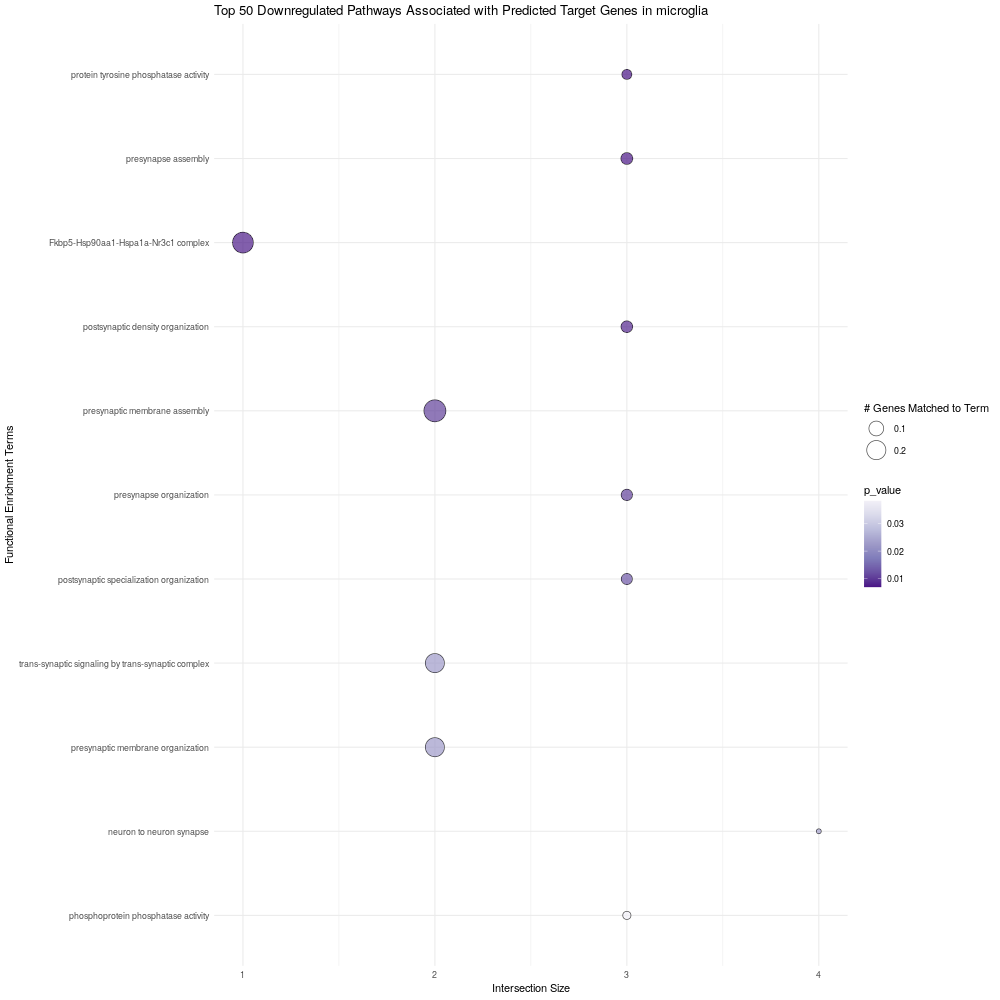

**Figure S9:** Dot plot of the top downregulated pathways (y-axis) associated with SETBP1 target gene set in microglia. The color of the dot corresponds to the significance (p-value; light purple to dark purple) and the size of the dot to the intersection size (proportion of genes matched to that term).

**Figure S10: gprofiler2 GO pathway analysis on predicted Setbp1 target genes upregulated in S858R microglia**
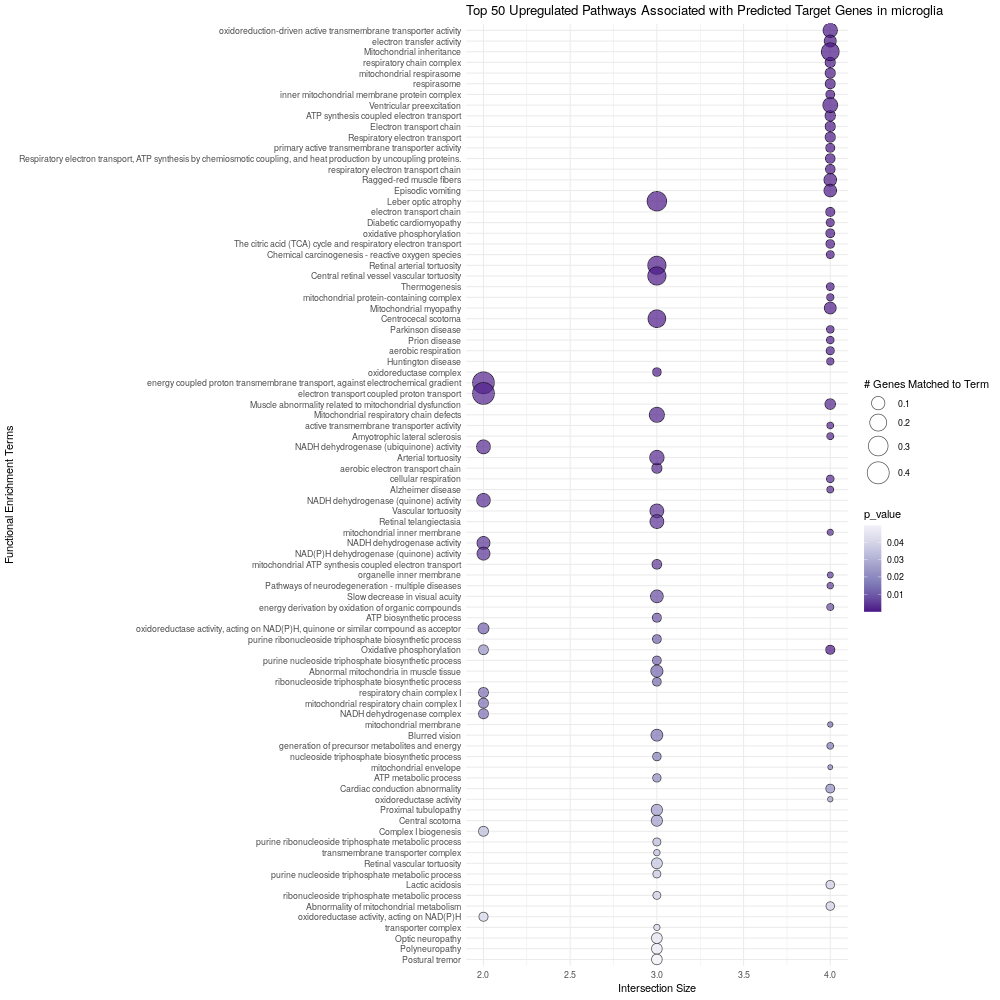

**Figure S10:** Dot plot of the top 50 upregulated pathways (y-axis) associated with SETBP1 target gene set in microglia. The color of the dot corresponds to the significance (p-value; light purple to dark purple) and the size of the dot to the intersection size (proportion of genes matched to that term).

**Figure S11: S858R kidney cystic kidney signature**

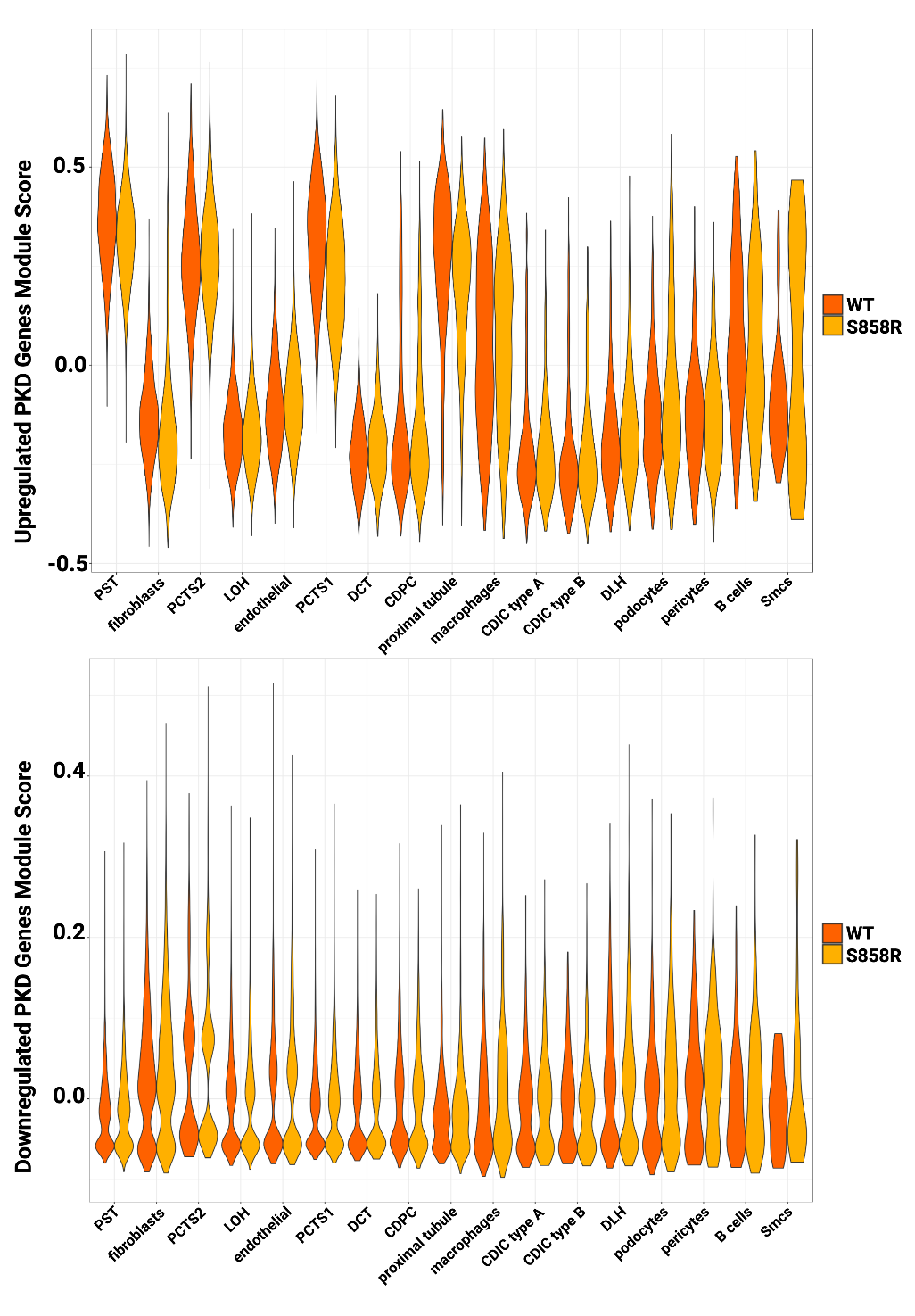
\

**Figure S11:** Split violin plot of module scores for PKD upregulated and downregulated gene signatures across kidney cell types in S858R and WT.

**Figure S12: S858R kidney failed repair signature**
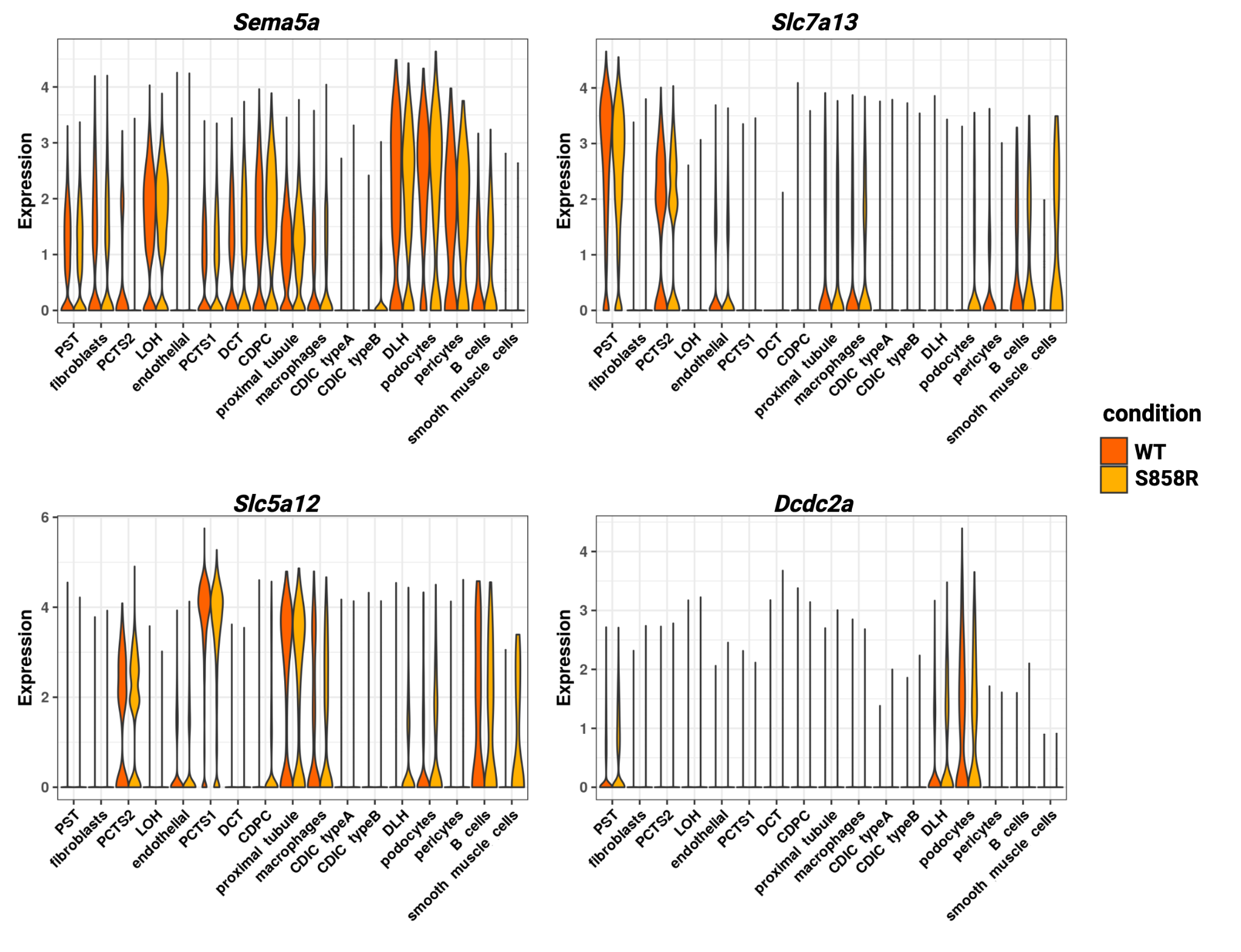

**Figure S12:** Split violin plot of expression of failed injury repair markers across kidney cell types in S858R and WT.

**Figure S13: SGS specific Hallmark pathways for cerebral cortex and all hallmarks for kidney with VISION analysis**

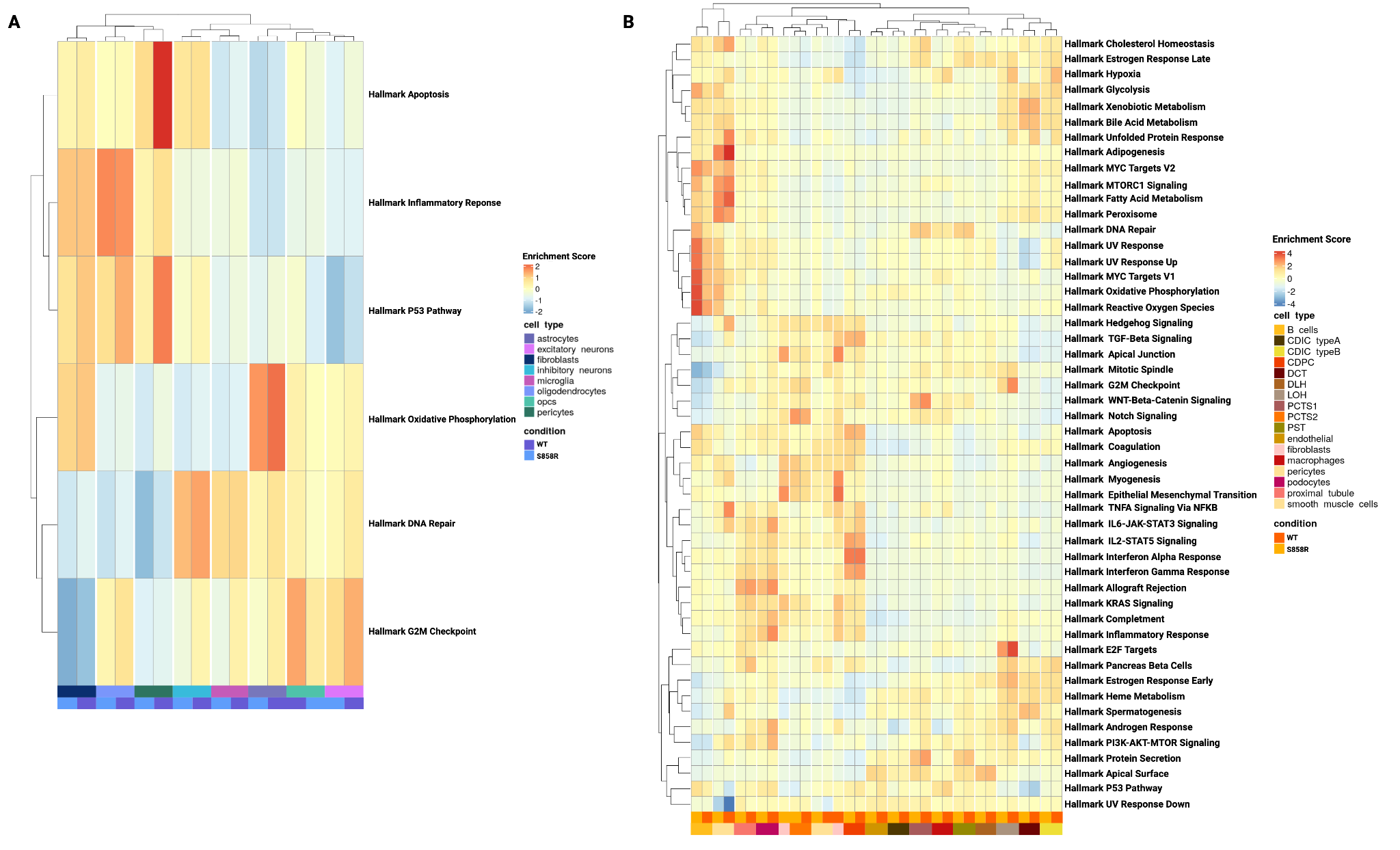

**Figure S13:** Heatmap showing enrichment of hallmark gene sets of Molecular Signatures Database (MSigDB) in each cell type of cerebral cortex **(A)** and kidney cortex **(B).**

**Figure S14: Expression of reactive astrocyte marker Gfap in cerebral cortex**

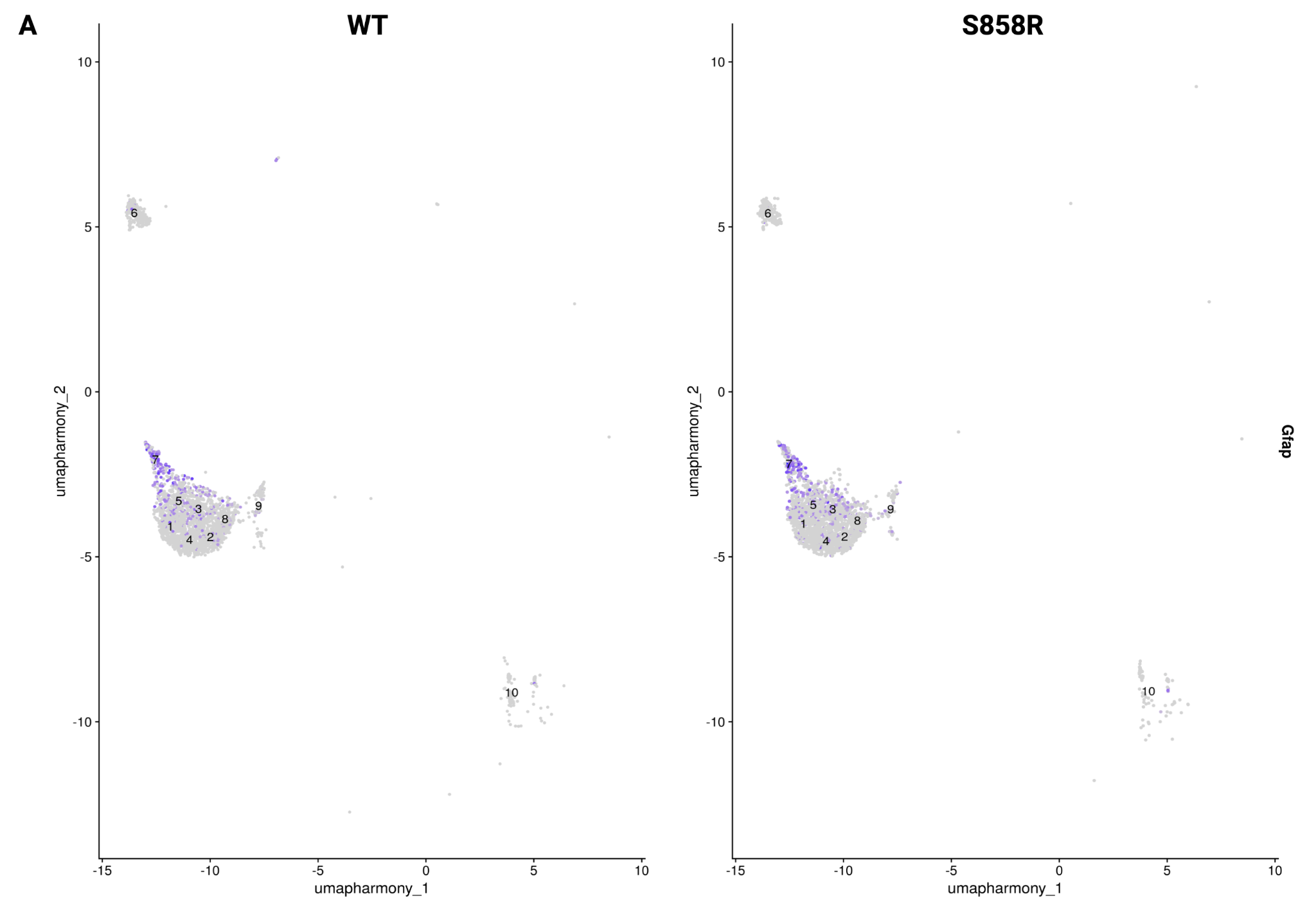

**Figure S14:** Expression of reactive astrocyte marker *Gfap* in astrocyte subclusters of WT (left) and S858R (right).

**Figure S15: Set has similar, ubiquitous expression across all cell types in both tissues**

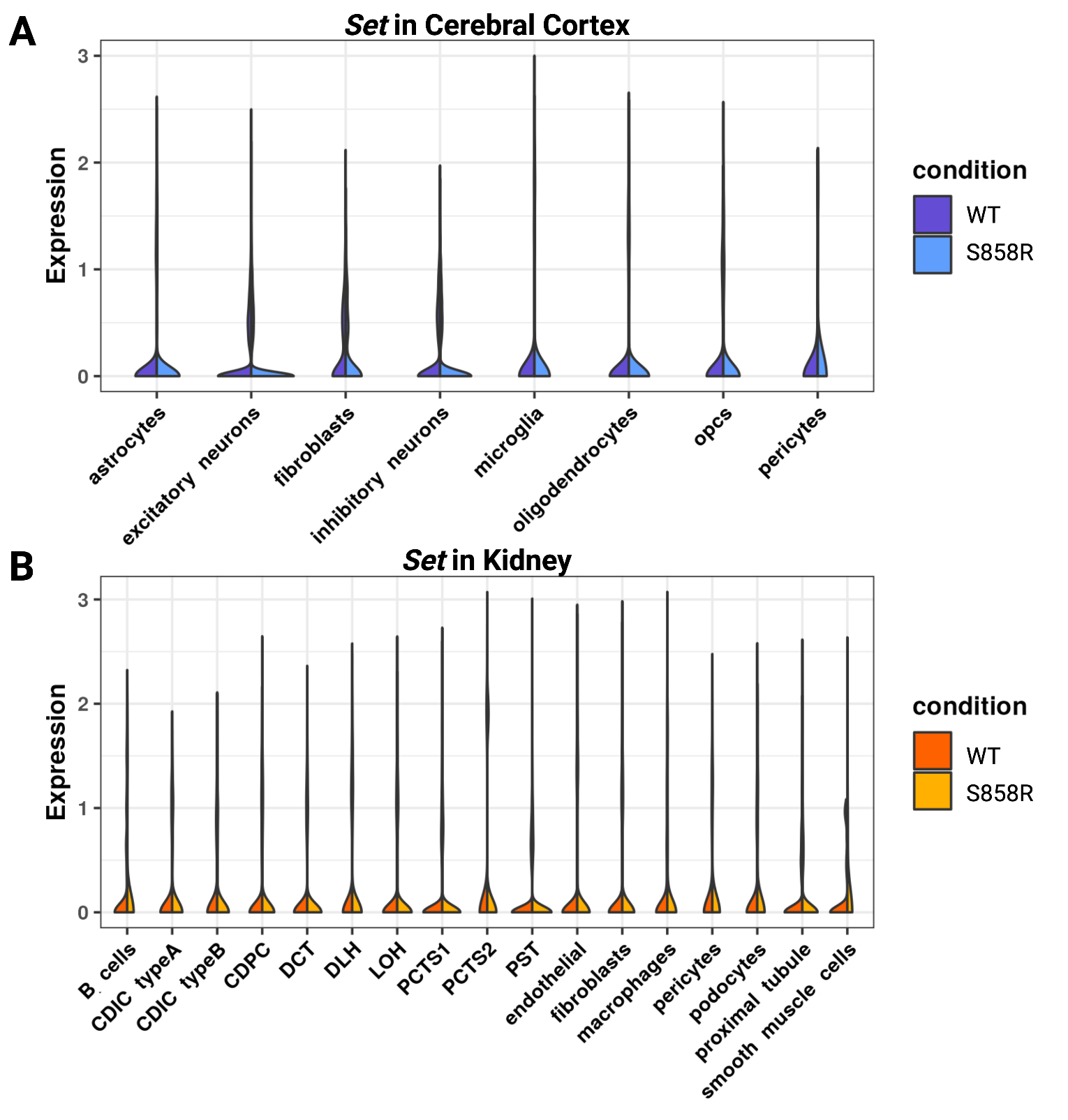

**Figure S15:** Split violin plot of expression of *Set* across cell types in both tissues, (A) cerebral cortex and (B) kidney in S858R and WT.

**Figure S16: Ppp2ca decrease in expression of S858R astrocytes (almost lack of) compared to WT whereas the inverse was observed in microglia and has similar expression across all cell types in kidney**

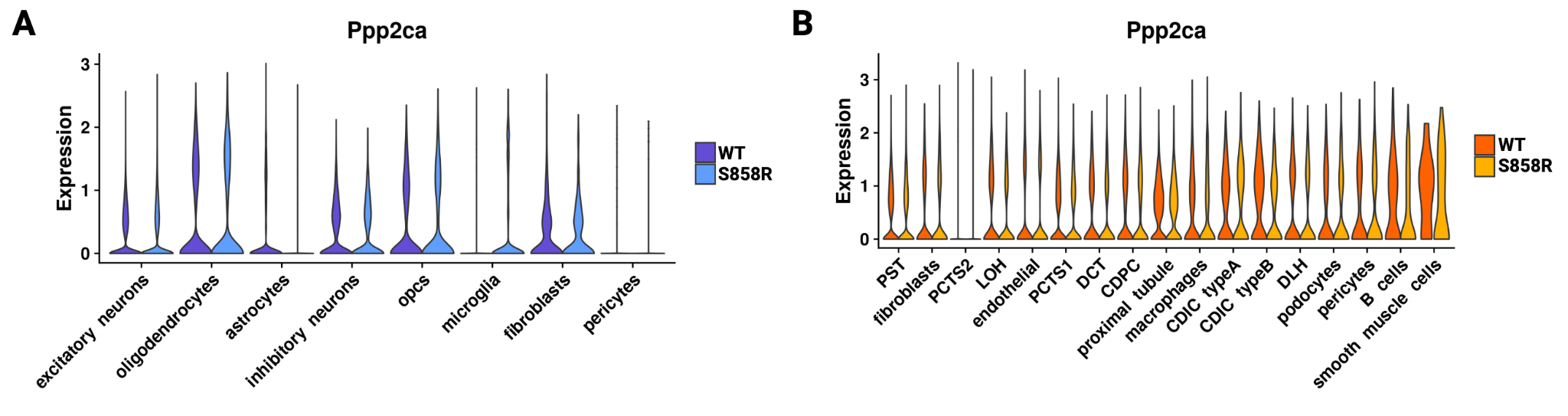

**Figure S16:** Expression of *Ppp2ca* in cerebral cortex (left) and kidney (right) for S858R and WT cell types.

**Table S7: Percentage and total number of differentially targeted genes by cell type for cerebral cortex and kidney**

Included in file Table_S7.xlsx

This table includes information for all cerebral cortex (blue) and kidney (orange) cell types and the percentage of TFs with TF targeting increased in the S858R or WT condition as well as the number of differentially targeted genes included by cell type.

**Figure S17: All Hallmark pathways for cerebral cortex with VISION analysis**

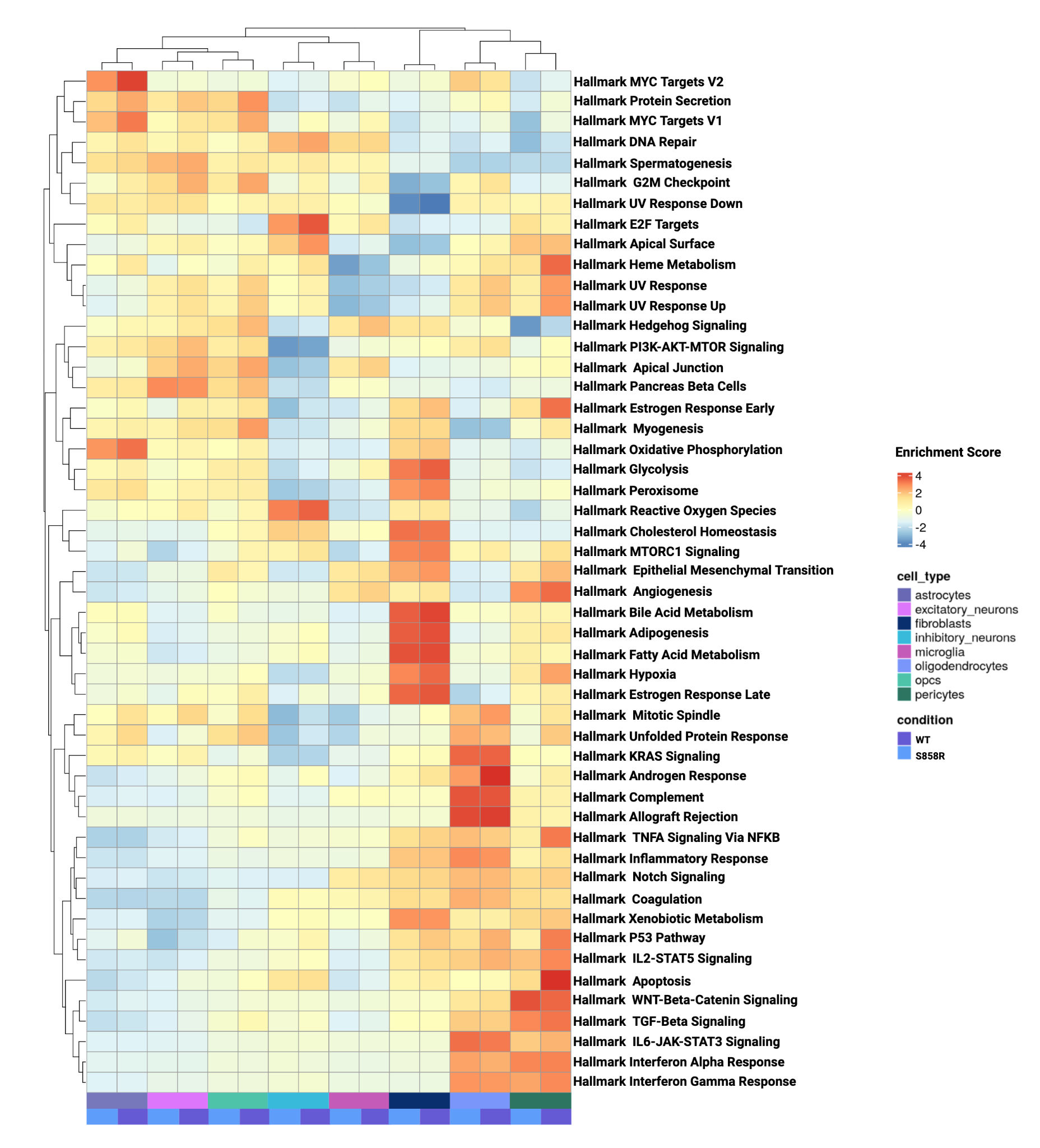

**Figure S17:** Heatmap showing enrichment of all hallmark gene sets of Molecular Signatures Database (MSigDB) in each cell type of cerebral cortex**.**

**Figure S18: Control Cooperativity and Regulation**
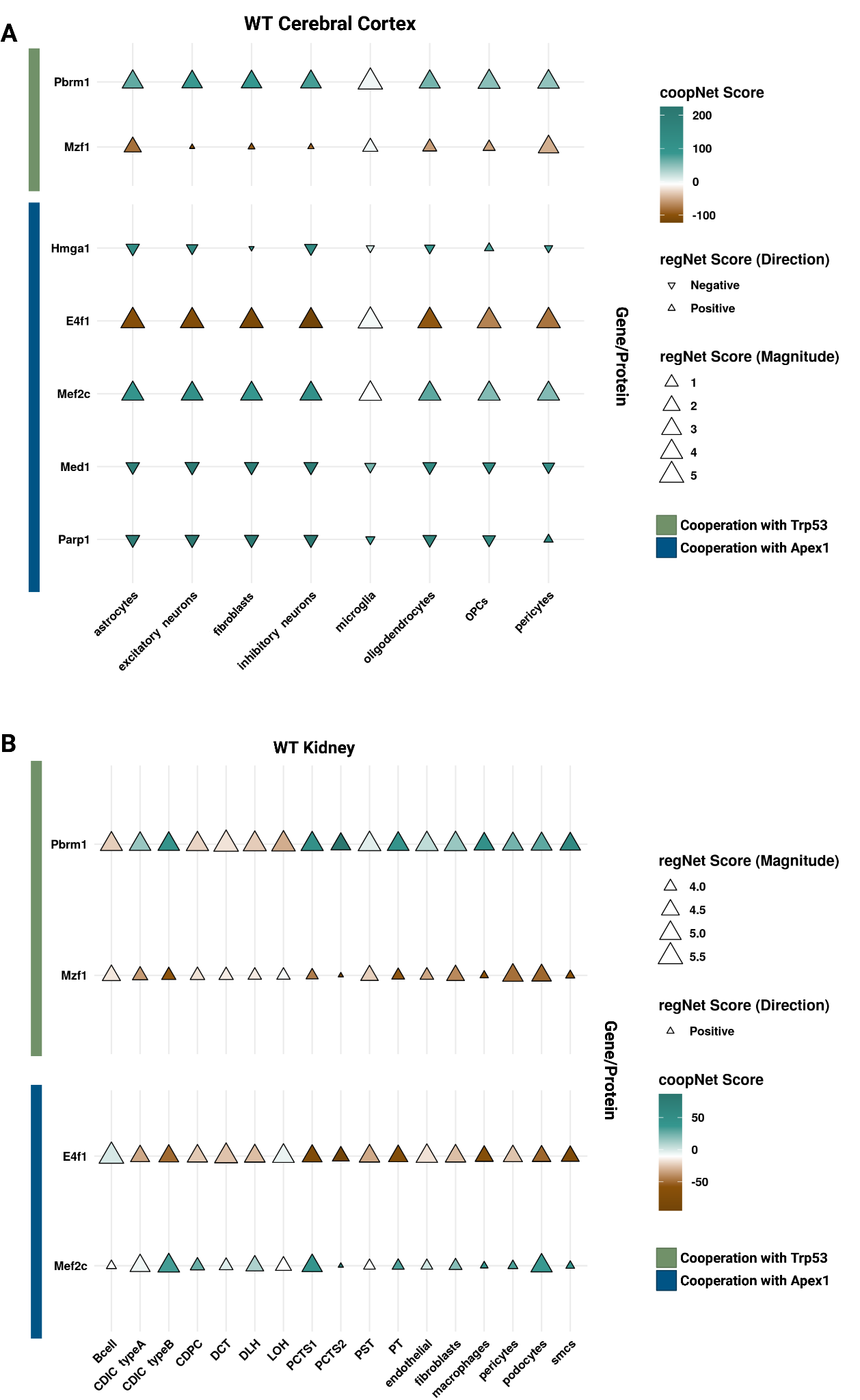

**Figure S18:**  Dot plot showing the Magnitude of regulatory (direction and size of triangle) and cooperativity (teal to brown) for network scores in the cerebral cortex **(A)** and kidney **(B)** of WT mice.

**Table S8: STRINGdb prior evidence for proteins of interest**

| **STRING score** | **Protein 1** | **Protein 2** |
| --- | --- | --- |
| 873 | Parp1 | Apex1 |
| 953 | Parp1 | Trp53 |
| 987 | E4f1 | Trp53 |
| 654 | Med1 | Trp53 |
| 207 | Apex1 | E4f1 |
| 735 | Trp53 | Pbrm1 |
| 287 | Apex1 | Hmga1 |
| 717 | Trp53 | Hmga1 |
| 396 | Trp53 | Mef2c |
| 537 | Trp53 | Mzf1 |

**Table S8:** Table of STRING scores representing as the strength of confidence in interactions between Protein 1 and Protein 2 from interactions identified in Figure 6.
